# A Keller-Segel model for *C elegans* L1 aggregation

**DOI:** 10.1101/2021.05.10.443398

**Authors:** Leon Avery, Brian Ingalls, Catherine Dumur, Alexander Artyukhin

**Affiliations:** Department of Applied Mathematics, University of Waterloo, Waterloo, Ontario, Canada; Department of Pathology, Virginia Commonwealth University, Richmond, VA, USA; Chemistry Department, State University of New York, College of Environmental Science and Forestry, Syracuse, NY, USA

## Abstract

We describe a mathematical model for the aggregation of starved first-stage *C elegans* larvae (L1s). We propose that starved L1s produce and respond chemotactically to two labile diffusible chemical signals, a short-range attractant and a longer range repellent. This model takes the mathematical form of three coupled partial differential equations, one that describes the movement of the worms and one for each of the chemical signals. Numerical solution of these equations produced a pattern of aggregates that resembled that of worm aggregates observed in experiments. We also describe the identification of a sensory receptor gene, *srh–2*, whose expression is induced under conditions that promote L1 aggregation. Worms whose *srh–2* gene has been knocked out form irregularly shaped aggregates. Our model suggests this phenotype may be explained by the mutant worms slowing their movement more quickly than the wild type.

**Author summary:** Among the most complex of animal behaviors are collective behaviors, in which animals interact with each other so as to produce large-scale organization. Starved first-stage larvae of the nematode *Caenorhabditis elegans* exhibit such a behavior: they come together to form aggregates of several hundred worms. How and why they do this are unknown. To address these questions, we developed a mathematical model of starved L1 aggregation. This model reproduced the main features of the behavior.

## Introduction

Among the most complex behaviors exhibited by the nematode *C elegans* are social behaviors such as mating [1] and aggregation. We recently described a new aggregation behavior in starved *C elegans* first-stage larvae (L1s) [2]. This new behavior raises two broad questions whose answers we lack: (1) How do starved L1s aggregate? I.e., what are the behavioral mechanisms by which they come together? (2) Why do starved L1s aggregate? What selective advantage (if any) do these mechanisms or aggregation itself provide? To aid in answering these questions, we describe here a simple mathematical model for L1 aggregation. In our first report of L1 aggregation behavior [2], we speculated on the answers to both question. This paper directly addresses only question (1).

L1 aggregation is not the only known *C elegans* aggregation behavior, and ours is not the first mathematical model of *C elegans* aggregation. It has been known for many years that in the presence of food (bacteria), most true wild isolates of *C elegans* aggregate, a behavior known as social feeding [3, 4]. Wild strains of *C elegans* prefer low concentrations of oxygen. The usual *C elegans* laboratory strain, N2, does not display social feeding at normal atmospheric oxygen pressure because of a gain-of-function mutation in the neuropeptide receptor gene *npr–1* [3]. We and others have speculated that the consumption of oxygen in an aggregate of worms lowers oxygen concentration and thereby attracts more worms [5], although this explanation is disputed [6]. Mathematical models of social feeding have recently been published [6, 7].

A third type of aggregation is mediated by indole-containing ascarosides [8]. L1s of *daf–22* mutants, which are unable to make ascarosides [9, 10, 11]) aggregated similarly to wild type [12]. Thus L1 aggregation is different from ascaroside-mediated aggregation. Observations of yet another type of aggregation have recently been published, together with a model [13]. This form occurs in the long-term survival form of the worm—the dauer larva—and is probably mediated largely by a simple physical mechanism, surface tension.

The model we present here is simpler than previous *C elegans* aggregation models in the following sense: it does not describe aggregation behavior in completely realistic detail. We attempt only to reproduce the essential aspects of the behavior. Accordingly, we simply assume the existence, which has been experimentally demonstrated [14, 15], of taxis mechanisms that allow worms to move in the direction they want to go. Although taxis mechanisms have been investigated for years, and much is known about them [e.g. 16–20], the model presented here is based on the idea that the end result of taxis (movement towards favored places) is sufficient to understand aggregation, and that mechanistic details are not essential.

A further simplification is to describe worms not as individuals, but via population density: a continuous function of space and time, *ρ*(*t*, **x**). We also propose a simple mechanism for interactions among worms via diffusible chemical signals. The resulting model takes the form of a system of partial differential equations (PDEs), a variation on the classic Keller-Segel [21] model, developed to explain the aggregation of cellular slime mold amoebae. Since its original publication, the Keller-Segel model has been the subject of much mathematical analysis [sections 11.1-11.3 of 22, 23, 24, 25]. Indeed, the Keller-Segel model, in its many variations, has become one of the classic models of pattern formation in mathematical biology. This model has the advantage of high mathematical tractability, both analytical [26] and numerical, the latter of which is the focus of this paper.

## Results

### Strategy

As described in the Introduction, our model is a deliberately simplified description of behavior that assumes the existence of taxis mechanisms that allow worms to move in the direction they want to go. Further, worms are modeled not as individuals, but as a continuous function of space and time, population density *ρ*(*t*, **x**). This simplification allows us to describe worm movement with a mathematically tractable partial differential equation (PDE) model similar to the well-studied Keller-Segel [21] model. Also, because worm density is a component of the model, it is straightforward to implement worm movement that explicitly depends on density. A disadvantage is that individual worms are not accurately represented by a continuous density function. Moreover, we ignore the fact that worms are worm-shaped: i.e., a worm is long and thin, with a head at one end and a tail at the other. Worm geometry is central to some other published models of *C elegans* aggregation [5, 6, 13].

We present two versions of the model: the precursor *attractant-only* model and the final *attractant+repellent* model. We begin with the simpler attractant-only model, which is closest in form to the original Keller-Segel model. This model was a partial success—it reproduced certain aspects of L1 aggregation seen in experiments, while failing in others. The more complicated attractant+repellent model better reproduced L1 aggregation behavior.

### Design of the PDEs

The attractant–only model for L1 aggregation consists of two coupled PDEs, a reaction-diffusion equation that describes the time evolution of the concentration of a diffusible chemical attractant, and a Fokker-Planck equation that describes the movement of the worms. The attractant PDE is:

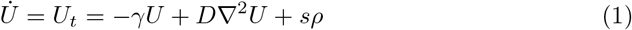

The worm PDE is:

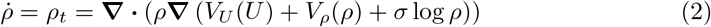

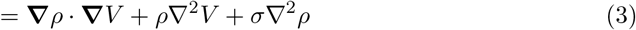

Functions and parameters appearing in (1 - 3) are listed in Table 1.

**Table 1.**
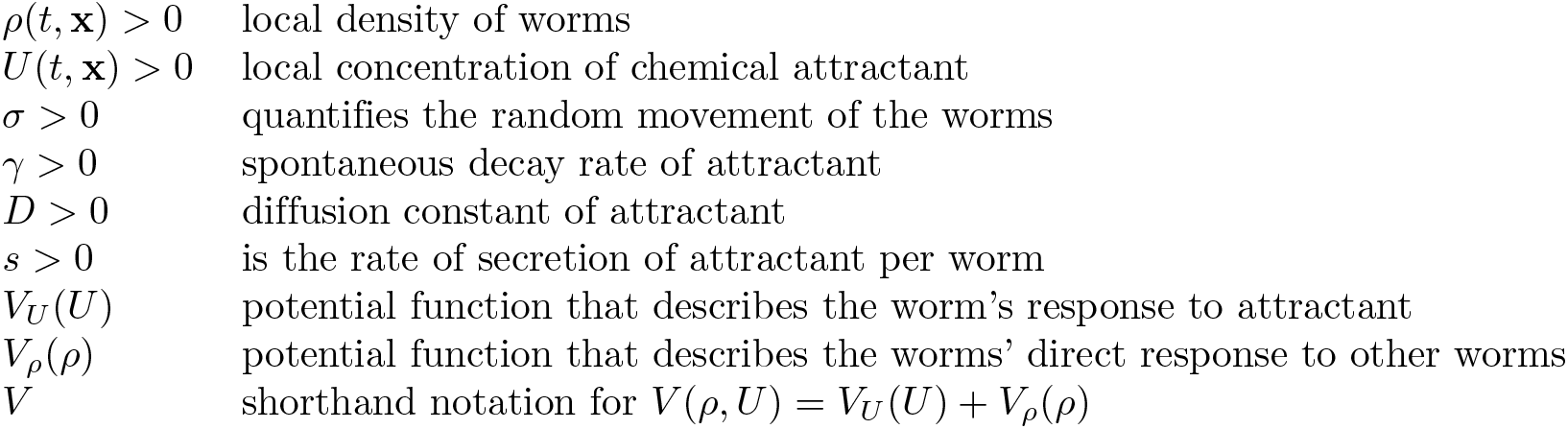
Functions and parameters appearing in the PDEs.

Intutively, the terms of (3) can be understood as follows. The first two terms describe how density changes when worms move towards lower potential. The first term, **∇***ρ* · **∇***V* arises from the movement of worms when the density is nonuniform. E.g., if density is low on the left and high on the right (**∇***ρ >* 0) and and worms in bulk are moving leftward (**∇***V >* 0) the density of worms at any fixed point will increase. The second term *ρ∇*^2^*V* describes increases in worm density when the worms converge towards a minimum of potential—∇^2^*V* is positive at and near minima. The final term, *σ∇*^2^*ρ* describes changes in density caused by random movement of the worms. Random movement tends to flatten out inequalities, so that density increases near minima of density (∇^2^*ρ >* 0) and decreases near maxima (∇^2^*ρ <* 0).

These equations are similar to those developed by Keller and Segel [21] to model the aggregation of *Dictyostelium discoideum* amoebae. Attractant PDE (1) is identical to the reaction-diffusion equation with which they model acrasin. Worm PDE (2) is a generalization of the equation they use to model the movement of amoebae, which, with specific choices of the potential functions *V*_*U*_ and *V*_*ρ*_, reduces to theirs.

In designing this model for L1 aggregation, we sought to reproduce certain general characteristics that were obvious in recordings of worm aggregation. First, the worms aggregate. This suggests that they are somehow attracted to each other. Given what we know about *C elegans* biology, it was an obvious guess that this attraction could be mediated by a diffusible chemical signal with limited range [15]. PDE (1) is essentially the simplest physically plausible that meets these criteria.

The design of the PDE describing the movement of the worms was more complicated. On the time scale of the experiments, neither birth nor death of new worms occurs. This suggested that it should be possible to express the rate of change of worm density as the divergence of some flow field. Since flux occurs by movement of worms, the net flux vector at any point is the density times the mean velocity of worms at that point. These considerations lead to a general equation of the form

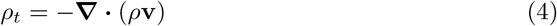

in which velocity **v** = **v**(*ρ, U*, **∇***ρ*, **∇***U*) is a vector field depending on density and attractant concentration and their gradients. We chose to assume that the velocity field is conservative, i.e. that it can be represented as the gradient of some scalar potential field. There is no compelling biological necessity for this assumption. We made it for two reasons: First it makes the PDE system more tractable analytically. Second, in recordings of worm behavior, we see that the worms eventually approach an equilibrium in which there is little net flow of animals, and no cyclic flows are obvious.

If the velocity field **v** is conservative, then it can be expressed as the negative of the gradient of some scalar potential field *V*. We chose a potential function that is a sum of a signal-dependent potential *V*_*U*_ and a density-dependent potential *V*_*ρ*_, for convenience in separately engineering signal and density dependence. This led to the final form (2).

For an attractant, *V*_*U*_ must be a decreasing function of signal. In early simulations with a linear *V*_*U*_ we encountered problems with numerical instability. Steep signal gradients frequently occur in the course of simulation. With a linear *V*_*U*_, these led to large velocities, which meant that worm density at one location was rapidly affected by density at distant locations. As a result it was impractical to satisfy the Courant–Friedrichs–Lewy (CFL) stability condition. We therefore sought a potential function whose dependence on *U* was convex.

A modification of Weber’s Law that closely corresponds to empirical data is

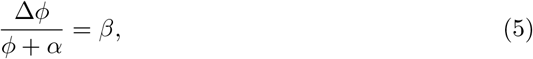

where Δ*ϕ* is the smallest detectable change in a stimulus *ϕ* and *α* and *β* are constants,. (This is Eq (1.2) of reference 27.) If we assume that just noticeable differences in stimulus represent equal changes in sensation *ψ*, we can integrate (5) to obtain the following psychophysical magnitude function,

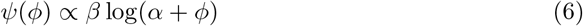

If attractant *U* is the stimulus and potential *V*_*U*_ the sensation, we get the following potential

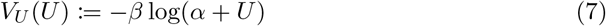

We negate *β* because worms move down a potential, and the potential for an attractant should thus be a decreasing function of its concentration. Parameter *β* determines the strength of attraction. The same potential with negative *β* describes a repellent.

We speculated that the circular shape of the aggregates [2] results from the worms packing together as tightly as possible. To reproduce this effect in simulations, we designed a density-dependent *V*_*ρ*_ potential that would reflect worms taking up space. The ideal would have been a hard sphere potential

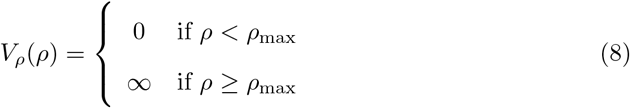

This potential function implies discontinuous time or spatial derivatives of density, and therefore functions poorly with numerical methods for solving the PDE system. We therefore approximated the discontinuity with a hyperbolic tangent function.

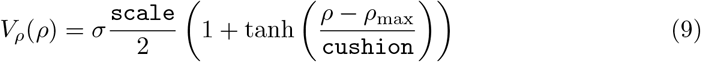

Four parameters determine the exact shape of *V*_*ρ*_: *σ, ρ*_max_, `scale`, and `cushion`. (We refer to the latter two by the symbols used to represent them in software code, since they will play little role in the mathematics.) Two parameters, *σ* and `scale`, determine the vertical scale. *σ* is the parameter that measures random worm movement (see (2)). *V*_*ρ*_ rises from near 0 for small values of *ρ* to *σ ×* `scale` for large *ρ*. Parameter *ρ*_max_ is the density at which *V*_*ρ*_ reaches half its maximum possible value. It is the point at which the *V*_*ρ*_ curve is steepest, and therefore the closest approximation to *ρ*_max_ of (8). Parameter `cushion` determines how abrupt the rise of *V*_*ρ*_ is. These functions are plotted in Fig 1. The Keller-Segel literature describes other, less-flexible, models in which organisms take up space [25], which we elected not to use.

**Figure 1.**
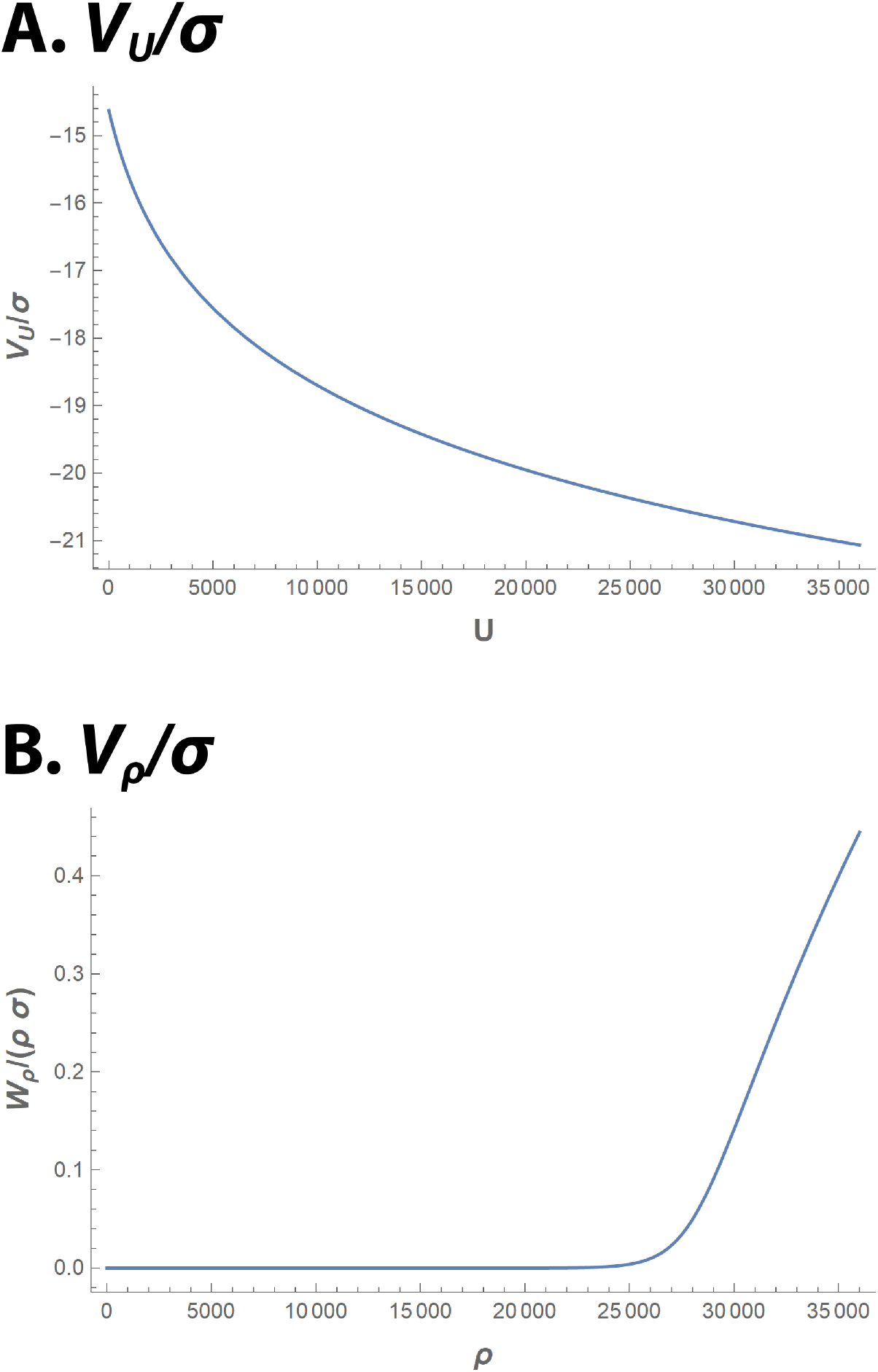
Potential function plots. Potential functions that appear in the *ρ* PDE (2). Both potentials are made dimensionless by dividing them by *σ*. Parameter values are as in Table 2.

### Parameter estimates

We required numerical estimates of parameters *γ, D*, and *s* that appear in (1) and *σ* of (2). In addition, we required values for *ρ*_max_, `scale, cushion` and *α* and *β* which determine the shapes of the potential functions *V*_*ρ*_ and *V*_*U*_.

**Table 2.** Parameter values.

|  |  |  |
| --- | --- | --- |
| $\bar{\rho}$ | mean worm density | $9000 \text{ cm}^{-\text{d}}$ |
| $\sigma$ | random worm movement | $5.555 \times 10^{-6} \text{ cm}^2 \text{ s}^{-1}$ |
| $\rho_{\text{max}}$ | midpoint of $V_\rho$ potential rise | $28\,000 \text{ cm}^{-\text{d}}$ |
| **cushion** | breadth of $V_\rho$ rise | $2000 \text{ cm}^{-\text{d}}$ |
| **scale** | height of $V_\rho$ rise | 2 |
| $\beta_a$ | strength of attraction | $1.111 \times 10^{-5} \text{ cm}^2 \text{ s}^{-1}$ |
| $\alpha_a$ | attractant concentration scale | $1500 \text{ cm}^{-\text{d}}$ |
| $\gamma_a$ | attractant decay rate | $0.01 \text{ s}^{-1}$ |
| $D_a$ | attractant diffusion constant | $1 \times 10^{-6} \text{ cm}^2 \text{ s}^{-1}$ |
| $s_a$ | attractant secretion rate | $0.01 \text{ cm}^{-\text{d}} \text{ s}^{-1}$ |
| $\beta_r$ | strength of repulsion | $-1.111 \times 10^{-5} \text{ cm}^2 \text{ s}^{-1}$ |
| $\alpha_r$ | repellent concentration scale | $1500 \text{ cm}^{-\text{d}}$ |
| $\gamma_r$ | repellent decay rate | $0.001 \text{ s}^{-1}$ |
| $D_r$ | repellent diffusion constant | $1 \times 10^{-5} \text{ cm}^2 \text{ s}^{-1}$ |
| $s_r$ | repellent secretion rate | $0.001 \text{ cm}^{-\text{d}} \text{ s}^{-1}$ |

A *C elegans* L1 is approximately a cylinder of diameter 15 µm and length 240 µm [28]. Since worms lie on their sides, a worm occupies approximate area 15 × 240 ≈ 3600 µm^2^. We chose the inverse of this area, 28 000 cm^−d^ as the parameter *ρ*_max_. (Here *d* = 1 or 2 is the spatial dimension. The same number, 28 000, was used for one and two-dimensional simulations to facilitate comparison.) Parameters `cushion` and `scale` have no real biological significance. Parameter cushion makes the ideal hard-sphere potential (8) continuous and differentiable, so that the PDEs can be solved numerically with differentiable functions. The value 2000 cm^−d^ worked. The `scale` parameter need only be chosen large enough to constrain the maximum density—we chose 2.

In small-scale simulations, we chose a mean density 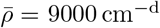 so that aggregates would occupy about 1*/*3 (i.e. 9000*/*28000) of the area.

Small molecules in water typically have diffusion constants in the range 1 × 10^−6^ cm^2^ s^−1^ to 1 × 10^−5^ cm^2^ s^−1^. (We assumed that the signals diffuse through the agar-solidified water under the worms. Worms also respond chemotactically to volatile chemicals diffusing through the air above them [15]—diffusion constants for such volatile signals would be much larger than for water-soluble signals.) We chose the diffusion constant of attractant, *D*_*a*_ =1 × 10^−6^ cm^2^ s^−1^, at the lower end of the range of diffusion constants in water. The aggregates that form have diameters of hundreds of micrometers. The mean distance a molecule of attractant diffuses before decaying is 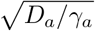. We therefore chose *γ*_*a*_, the decay rate of attractant, to give it a range 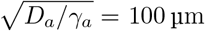. To fulfil its role in the model, the repellent, introduced below, needs to have a longer range, so we chose a large diffusion constant of *D*_*r*_ =1 × 10^−5^ cm^2^ s^−1^ and a smaller decay rate, giving it a range of 1 mm.

Parameters *s*_*a*_ and *s*_*r*_, the rates at which a worm secretes attractant and repellent, effectively set the units of concentration. We chose units of concentration such that *s*_*i*_ and *γ*_*i*_ (for *i* = *a* or *r*) were numerically equal. (That is, if concentration is measured in “number of units of stuff”/cm^d^, we chose the units in which “stuff” is measured to be the amount secreted by one worm in one mean lifetime of the stuff, i.e. 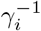. This has the effect that if *γ*_*i*_ = ⟨number ⟩ s^−1^, then *s*_*i*_ = ⟨number⟩ “stuff units” cm^−d^s^−1^, with the number being the same in the two cases.) This ensures that concentrations *U*_*i*_ and worm density *ρ* are in the same range numerically.

Artyukhin et al found that the minimum worm density for aggregation is 1500 cm^−2^ [2]. We identified this with the density threshold for instability. We chose *α*_*a*_ = *α*_*r*_ = 1500 cm^−2^, to make *V*_*U*_ linear near the threshold, and to be obviously convex near *ρ*_max_. We then chose *β*_*a*_ = 2*σ* to reproduce the 1500 cm^−2^ density threshold for instability in the attractant-only model. For the attractant+repellent model described below, we kept this value for *β*_*a*_ and chose *β*_*r*_ = −2*σ* for the repellent. Adding repellent to the model increased the calculated instability threshold to 2357 cm^−d^.

Parameter *σ* determines how rapidly the worms spread. Artyukhin et al [2] found that worms placed at the center of a 6 cm diameter petri plate spread to occupy much of the area of the plate in 12 h, but at this time they still remain mostly concentrated near the center. To estimate *σ*, we asked what value of *σ* would reproduce the observed behavior of worms in circular petri plates.

We began by choosing values of *σ* that would approximately reproduce this distribution if the worms’ motions were purely diffusional, i.e., if their motions were governed by

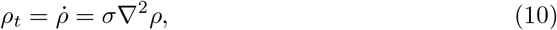

with Neumann boundary condition

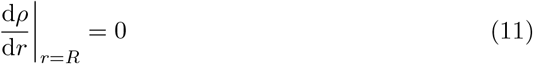

Here *R* = 3 cm is the radius of the petri plate. Eqs. (10, 11) can be solved by separation of variables. Any solution *ρ*(*t, r, θ*) can be represented as a sum of exponentially decaying eigenfunctions of the Laplacian. The circularly symmetric eigenfunctions on the disk with Neumann boundary condition (11) are

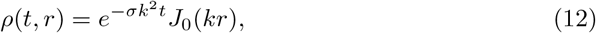

where *J*_0_ is a Bessel function of the first kind. The wavenumber *k* must be chosen to satisfy boundary condition (11), i.e. *k* = *j*_1,*n*_*/R*, where *j*_1,*n*_ is the *n*^th^ nontrivial zero of *J*_1_. The smallest wavenumber, corresponding to the circularly symmetric eigenfunction that decays most slowly, is thus *k* = *j*_1,1_*/R* ≈ 3.8317*/*3 cm ≈ 1.28 cm^−1^. We began by choosing *σ* so that the corresponding time constant 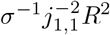 was approximately 12 h. Of course, the motion of the worms is not purely diffusional. We therefore refined our estimate of *σ* by numerical solution of the attractant+repellent system described below, from an initial condition in which the worms began near the center of the petri plate. From these simulations we chose *σ* =5.555 × 10^−6^ cm^2^s^−1^ as producing results that resembled experimental results.

Table 2 summarizes parameter values.

### Simulation results, attractant-only model

Fig 2 shows results of numerical simulations of the attractant-only model.

**Figure 2.**
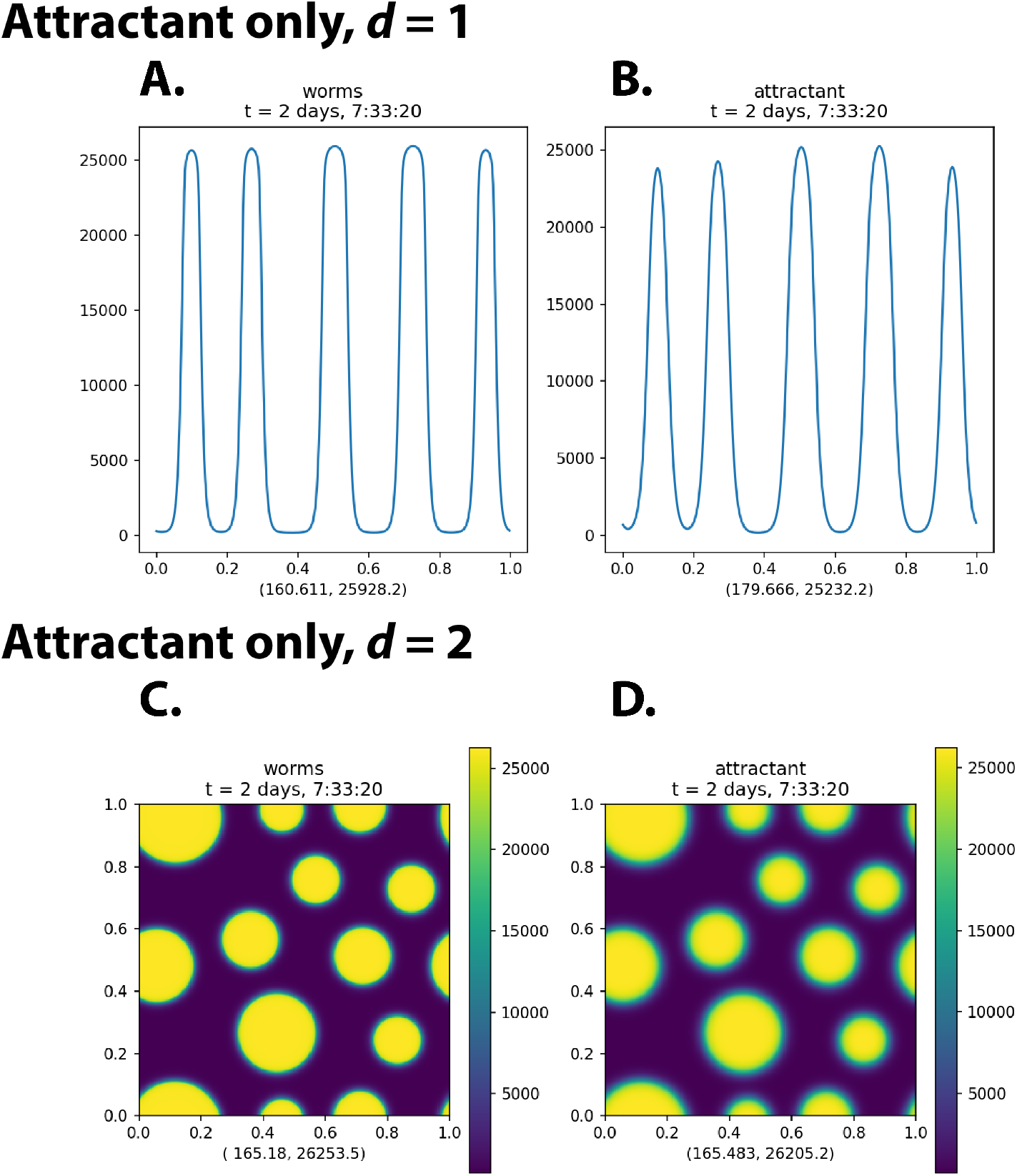
Simulation of the attractant-only model. This figure shows the state of a numerical simulation of the attractant-only model after 200 000 s (2 days and 7 hours). The initial condition was a uniform worm density of 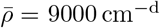, perturbed by normally distributed random noise of standard deviation 1% (i.e. 90 cm^−d^). (The entire time courses can be seen in the videos `options138a.mp4` and `options139.mp4` in the Supporting Information.) Supporting Information Fig S1 shows results at *t* = 200 000 s and *t* = 1 ×10^7^ s (116 days) of ten independent runs of the same simulation with different pseudorandom noise in the initial condition. Panels **A, B** show the results of simulations in one-dimensional space; **C, D** show results in two-dimensional space. **A, C** show density *ρ*; **B, D** show attractant concentration *U*. The two numbers below each plot are the minimum and maximum values of the plotted function over the entire 1 cm × 1 cm domain. The spatial units are centimeters.

This model successfully reproduced the experimental results in certain ways, but failed in others. It was successful in that circular aggregates of maximum density rapidly formed. The aggregates had sharp boundaries, and outside of aggregates worm density was low and uniform. This is most easily seen in the one-dimensional results (Fig 2A), but is also true in two dimensions. These are also characteristics of the experimental results. (See Supporting Information video `N2_5e5_washed.avi`.)

The model failed to reproduce the patterning of aggregates. In experiments (Supporting Information video `N2_5e5_washed.avi`), aggregation appears to reach an equilibrium after 12 h. Although individual worms continue to move actively, the aggregates themselves show little change after the first several hours. Those aggregates that form after the worms disperse from where they are initially placed are never larger than ca. 700 µm in diameter. Most of the worms, in fact, end up in aggregates close to this maximum size. These aggregates are also fairly uniformly spaced—the distance from one aggregate to its nearest neighbors varies little.

In numerical solutions of the attractant-only model, however, aggregates had no maximum size (aside from that imposed by the fixed finite number of worms), and their spacing was not uniform. Furthermore, even after 200 000 s, they were not at equilibrium. This can be seen by computing the velocity *v* = ‖**∇***V* ‖, which at equilibrium would be zero everywhere, but remained well above zero throughout the simulation. More obviously, it is seen by continuing the solution past *t* = 200 000 s. Fig S1 shows that aggregates increased in size and decreased in number between *t* = 200 000 s and *t* = 1 × 10^7^ s. In fact, we believe the only true equilibria of the attractant-only model are those in which there is a single large aggregate containing almost all the worms. This state was reached at *t* = 1 × 10^7^ s in one of the ten simulations in Fig S1.

In fact, this observation is consistent with linear stability analysis of the attractant-only model (see Supplemental Information). PDE system (1, 2) shares with the original Keller-Segel system the property of density-dependent instability. The condition for a sinusoidal variation of wavenumber *k* to be unstable is (35). The condition for instability (35) has no minimum wavenumber other than zero. Wavenumber is inversely proportional to wavelength, so that there is no nonzero minimum wavenumber means there is no natural maximum size for the aggregates that form when the density exceeds threshold. (This is a well-known property of the classical Keller-Segel model as well—see, for instance, section 11.3 of reference 22.) It is true that the attractant has a natural range, 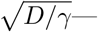the distance an average molecule diffuses before it decays. However, worms attract each other, albeit weakly, even when they are further apart than this. There is thus no mechanism in the model (1, 2) that would prevent the merging of aggregates to unlimited size. This is true for any attractant-only Keller-Segel model.

### A repellent is necessary

We could not reproduce the experimental observed uniformity of aggregate size in numerical experiments with attractant-only models. We suspected that the addition of a negative signal to oppose the attractant, a repellent, would solve the scale problem. Linear stability analysis supports this intuition. (See Supporting information section Linear stability analysis of the attractant+repellent model.)

Intuitively, what one requires is a short-range attractant and a long-range repellent. We therefore added to the attractant-only model a repellent with diffusion constant *D*_*r*_ = 10*D*_*a*_ and decay rate *γ*_*r*_ = 0.1*γ*_*a*_. The range of this repellent 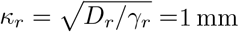 is ten times that of attractant, and is approximately equal to the observed spacing between aggregates. Thus, in the attractant+repellent model, attractant PDE (1) is replaced with two PDEs, one (13) for attractant and the other (14) for repellent.

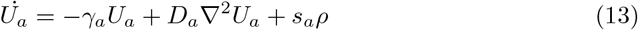

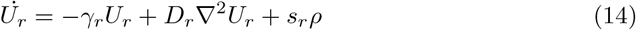

As shown in Fig 3, addition of a repellent to the model produced the predicted effect. Aggregates formed with characteristic size and spacing approximately matching those seen in experiments on worms. These solutions are close to equilibrium at *t* = 200 000 s, as seen by comparison with the results at *t* = 1 × 10^7^ s. (Compare Figs S2A and B.) There is even a hint of pattern formation, with the aggregates in an approximate hexagonal array at *t* = 200 000 s. The hexagonal patterning is near perfect at *t* = 1 × 10^7^ s, with the exception of lattice defects, most easily recognized as slightly smaller aggregates surrounded by five rather than six neighbors. (Fourier analysis confirms the regularity of these patterns [26].) If there is a problem with the result, it is that the array is too perfect. More irregularity is seen in experiments with real animals.

**Figure 3.**
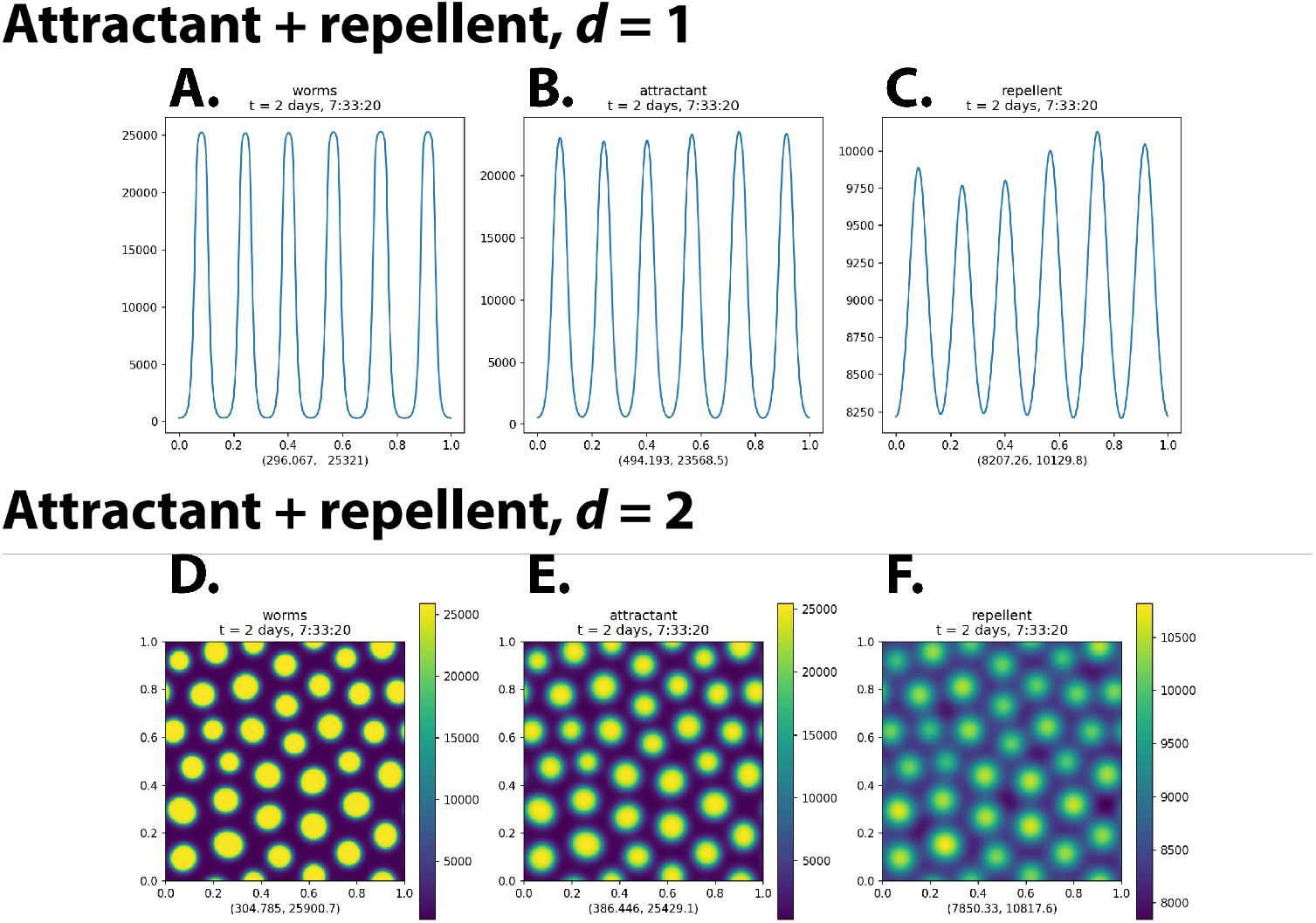
Attractant+repellent simulation Panels **A, D** show density *ρ*, **B, E** show attractant concentration *U*_*a*_, and **C, F** show repellent concentration *U*_*r*_. The spatial units are centimeters. The two numbers below each plot are the minimum and maximum values of the plotted function over the entire 1 cm ×1 cm domain. Note the different scale of the attractant and repellent plots. The means are the same, but because repellent is a longer-range signal, it is smoothed much more by diffusion and varies less than attractant. (Supporting Information videos `options140a.mp4` and `options141.mp4` show the full time courses for these simulations. Supporting Information Fig S2 shows ten independent solutions of the two-dimensional system with different pseudorandom noise at time 0.)

### *srh–2* encodes a G–protein coupled receptor expressed in starving L1s

Attempting to understand molecular mechanisms of L1 aggregation, we measured gene expression in starved L1s in the presence and absence of ethanol or acetate, either of which is required for aggregation [2]. We identified an ethanol-induced gene, *srh–2*, whose expression increases at the time that starved L1s become capable of aggregation (Supporting information). Gene *srh–2* is predicted to encode a sensory receptor, i.e., a protein expressed on the surface of a sensory neuron, capable of detecting chemicals in the environment. To find out whether *srh–2* plays a role in L1 aggregation, we knocked the gene out. (That is, we genetically engineered a mutant strain that lacks a functional *srh–2* gene.) We then tested the *srh–2* knockout worms for aggregation. As shown in Fig 4, these mutant worms still aggregate, but the aggregates are irregular in shape. Furthermore, the number of worms outside large aggregates is larger in *srh–2* than in wild-type. In Fig 4B,C one can see that the frequency of isolated individuals and of small aggregates are both elevated.

**Figure 4.**
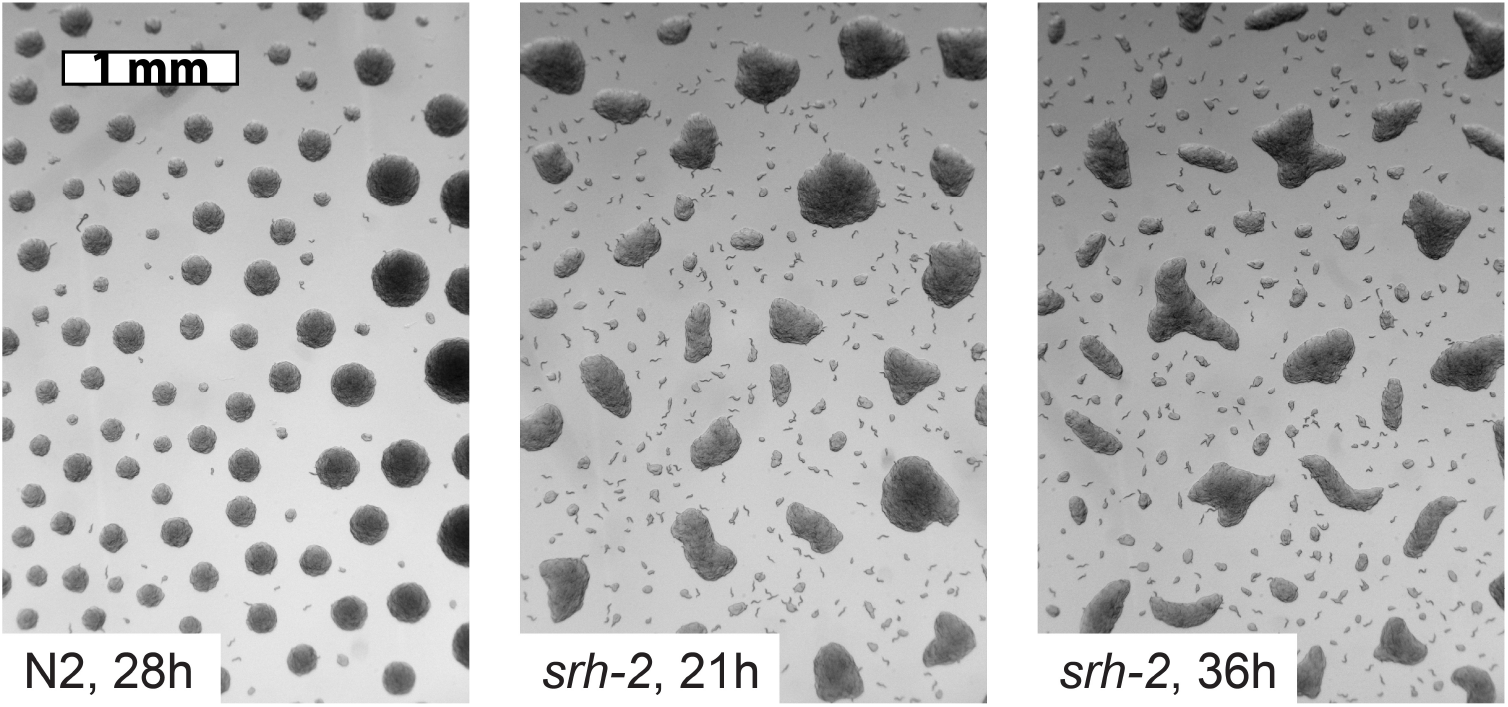
*srh–2* knockout L1 aggregation Starved L1s of mutant worms lacking a functional *srh–2* gene aggregate, but the aggregates they form are irregularly shaped (the *animal crackers* phenotype).

### The Srh–2 phenotype may be modeled by rapid decay of worm movement

Two observations suggested a partial explanation of the *srh–2* phenotype. First, in the movies of the attractant+repellent simulation, one sees formation of irregularly shaped aggregates at early times. With time, these aggregates become circular. Second, in movies of aggregating L1s, there is a lot of rapid movement at early times, but as time goes on, fewer worms are seen moving. This suggested that worm movement might slow with time, perhaps because the worms run low on energy. (They are, after all, starving.)

Together, these observations suggested an explanation for the Srh–2 phenotype–—perhaps the movement of *srh–2* knockout worms slows down faster. In the model, such a movement slowdown would be reflected in the decrease of the parameters *σ* (representing random worm movement) and *β*_*a*_, *β*_*r*_ (representing signal–responsive movement) with time. We modeled slowdown with the attractant+repellent PDEs (2, 13, 14), but with parameters *σ, β*_*a*_, *β*_*r*_ time-dependent:

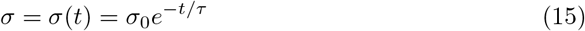

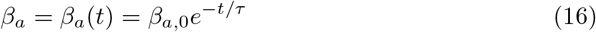

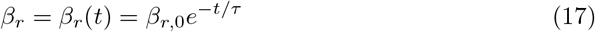

(Note that this is not the same as simply stretching the time axis of the attractant+repellent model, because the time–scales of Eqs (13, 14) remain unchanged.) The *t* = 0 values *σ*(0), *β*_*a*_(0), *β*_*r*_(0) were the same as those of *σ, β*_*a*_, *β*_*r*_ in Table 2. Fig 5 shows results at *t* = 200 000 s of simulations of the slowdown model with four different values of *τ*. When *τ* is very small (e.g. 30 min, Fig 5A) aggregation is arrested before dense aggregates form. Larger values of *τ* permit the formation and persistence of irregular aggregates. For small enough values of *τ* (Fig 5A,B), we also see an elevated background worm density.

**Figure 5.**
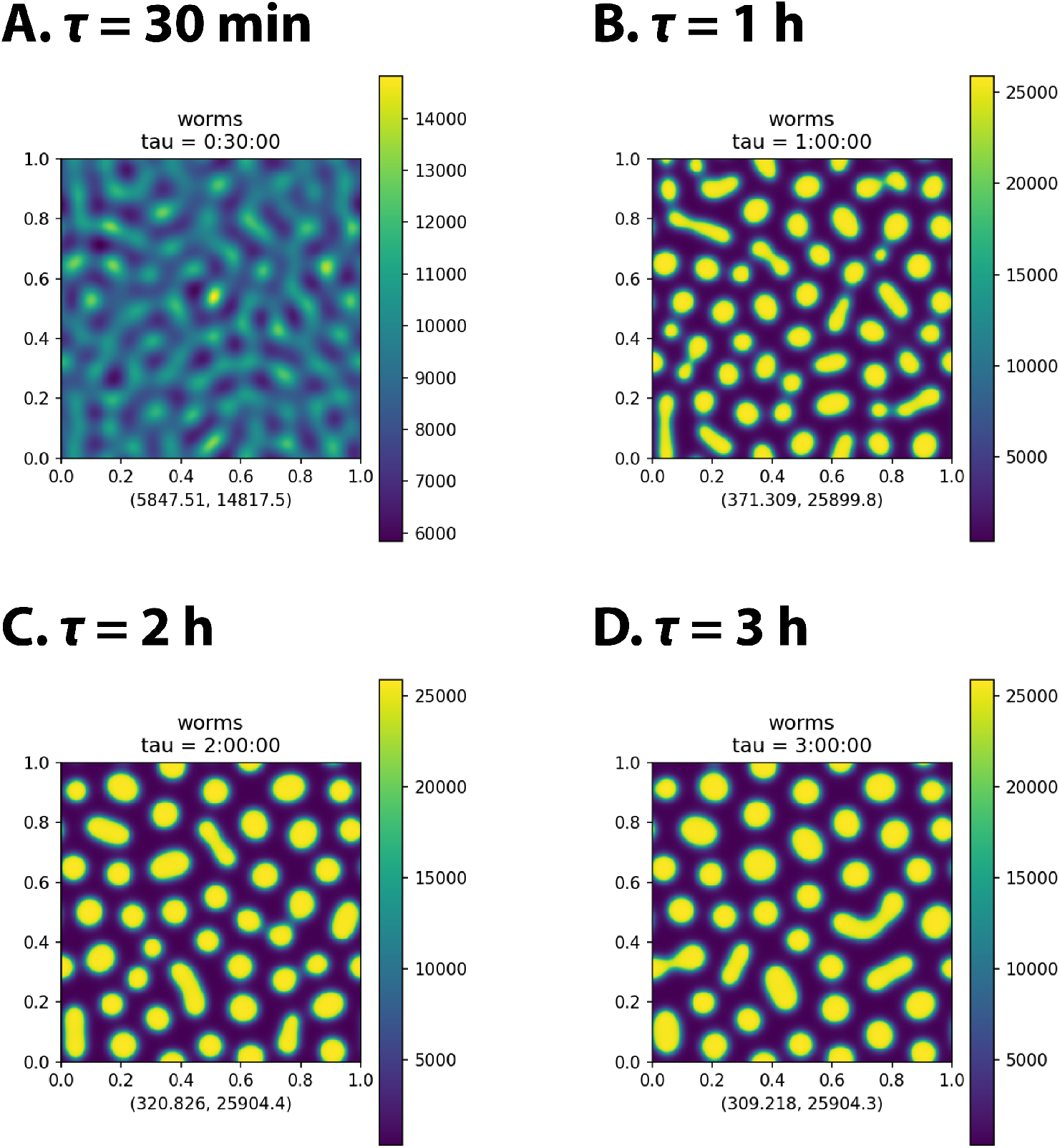
Attractant+repellent simulation with slowdown Worm density *ρ*(*t*, **x**) of a slowdown model simulation at *t* = 200 000 s for four different values of *τ*.

### Individual-based simulations

To check our PDE model, we also simulated a cellular Potts individual-based model in the Morpheus [29] modeling environment (details in Methods section). Fig S3 shows result that can be compared to Figs 2C,D and 3D,E,F. The results are similar. It has not yet been computationally feasible to reproduce Figs S1, S2, and 6. (These computations are in progress.)

### Full-scale simulations

Our experimental studies of aggregation usually begin by placing a large number of worms in the center of a 6 cm petri plate [12]. (See Supporting video `N2_5e5_washed.avi` which records the behavior over 12 h of 500 000 worms that were placed on the center of a plate at time 0.) These experiments begin with the dispersal of the worms, so that the density is high near the center of the plate and lower towards the edges. To more closely mimic such experiments, we solved the attractant+repellent model on a 6 cm × 6 cm square, from an initial condition in which most of the worms began near the center of the plate. Fig 6 shows the distribution of worms at *t* = 200 000 s.

**Figure 6.**
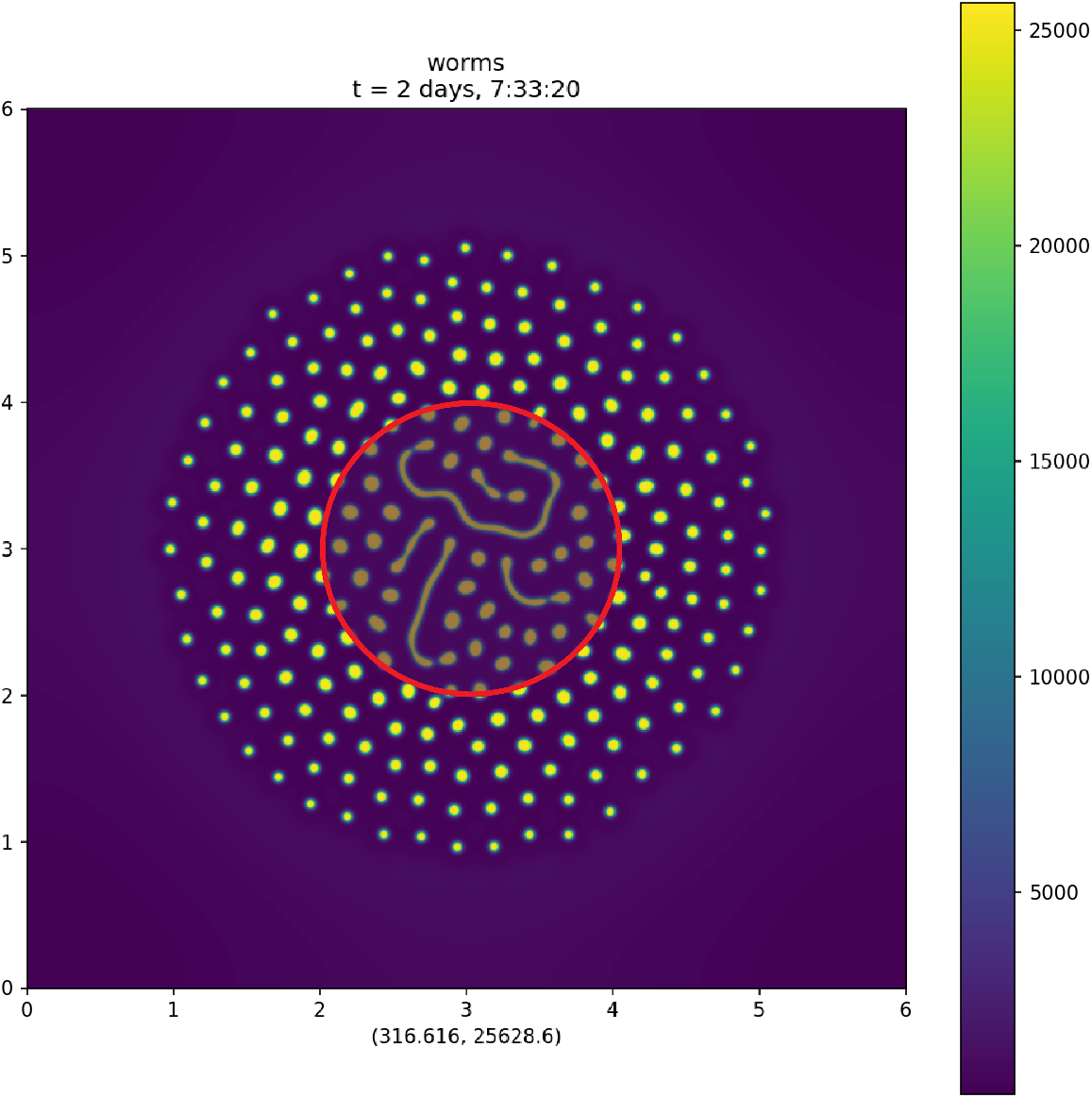
Full-scale attractant+repellent simulation Simulation of the attractant+repellent model on a 6 cm × 6 cm domain. The simulation began with 68 400 worms in a 2 cm diameter circle at the center of the plate (inner red circle). (Supporting Information video `options157.mp4` shows the entire time course.) The spatial units are centimeters. The two numbers below the plot are the minimum and maximum values of the plotted function over the entire 6 cm × 6 cm domain. Density is muted in a central 2 cm diameter circle, corresponding to where the worms were initially placed, to suggest the region in which we think influences not included in our model might be important.

We do not believe that the attractant+repellent model accurately represents the physics and biology of worm motions in the region near the center of the plate. For instance, in the preparation of eggs from which the L1s used for the experiment hatch, some non-living debris is inevitably generated. This debris, which is transferred to the center of the plate along with the worms, may influence behavior.

Outside this central region, the behavior of the full-scale simulation resembled the behavior of worms on a 6 cm diameter petri plate.

### Spectral comparison of experimental and simulation results

As a first approach to quantitative evaluation of the similarity of simulation results to experimental, we compared Fourier Power spectra of the final image of experimental video `N2_5e5_washed.avi` to the 200 000 s density function from full-scale simulation (Fig 7). (See the Methods section below for details.) Two-dimensional spectra (Fig 7C,D) show prominent rings at wavenumber *k* = ‖**k**‖ ≈ 10 cm^−1^, showing the existence of periodic structure with wavelength approximately 1 mm. The approximate circular symmetry of the power spectra result from the approximate circular symmetry of the images.

**Figure 7.**
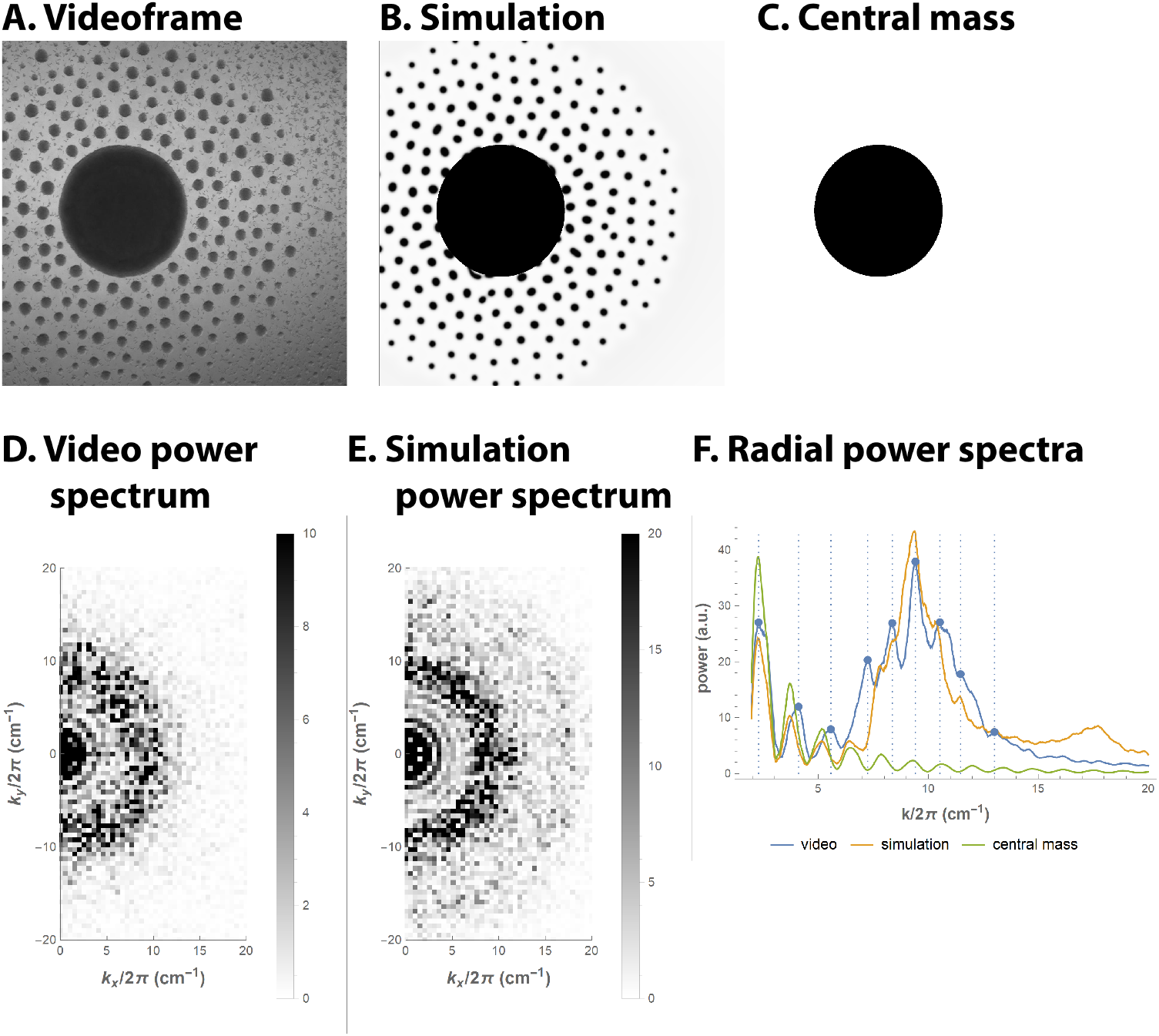
Spectral comparison of experimental and simulation results **A**. Last frame of aggregation video N2 5e5 washed.avi, cropped to a square to facilitate Fourier analysis. This square is 1.93 cm in size. **B**. Final time point of the full-scale simulation 6, cropped and scaled to match **A** as closely as possible, with a corresponding central mass added. **C**. The central mass alone. **D**. Low-frequency region of the two-dimensional power spectrum of the discrete Fourier transform of image **A**. The power scale is truncated at 10 AU (arbitrary units) **E**. Low-frequency region of the power spectrum of image **B**. The power scale is truncated at 20 AU. **F**. Radially summed power spectra of images **A**,**B**, and **C**. Peaks of the experimental image spectrum are picked out in blue.

To obtain higher resolution spectra, we computed radial power spectra by summing power *p*_**k**_ for spectral components with equal or approximately equal *k* = ‖**k**‖. These radial spectra (Fig 7F) show additional substructure within the *k* ≈ 10 cm^−1^ ring. The radial spectrum of the experimental image has about six peaks between 7 and 13 cm^−1^.

The simulation spectrum has peaks or shoulders at the locations of some but not all of the experimental spectrum peaks. (Note that we adjusted the scaling of the simulation image to make the largest peaks near 9.5 cm^−1^ match, so the coincidence of this particular peak is not significant.)

In both Fig 7E and F a weak feature is visible in simulation results at about 18 cm^−1^. Rather than suggesting structures with wavelength 0.5 mm, this may be a harmonic of the 1 mm periodicity. No such feature is visible in experimental results. It may be that the experimental data are too noisy for such small structures to survive in the power spectrum.

## Discussion

### Summary

Our final model, the attractant+repellent model, appeared to reproduce the main features of L1 aggregation. Spectral analysis suggests that the spatial patterning of aggregates in simulations resembles experimental results, although further work along these lines will be necessary. This model is minimal, we believe, in the sense that no simpler model of the Keller-Segel form can adequately reproduce the experimentally observed behavior. In particular, the observed patterning of aggregates—their roughly uniform spacing and sizes, required a short-range attractive influence and a longer-range repulsive influence. Luca et. al. [30] similarly conclude that a two-signal model is necessary to reproduce the patterning of senile plaques in their Keller-Segel model of Alzheimer disease.

In our model the attractive and repulsive influences took the form of labile diffusible small molecules produced by the worms. Some other possibilities that can be imagined work equally well. For instance, the attractant could be replaced by a ubiquitous repellent that is locally destroyed by the worms. (In some explanations of social feeding, oxygen plays this role.) It is even possible that the attractant and repellent are the same molecule, if it has the unusual behavioral characteristic of being repulsive at long range (i.e., lower concentration) and attractive at short range (high concentration). Either the attractant or the repellent could be replaced with a physical force, e.g. the physical attraction produced by surface tension (which, however, has too short a range to work well in our current models).

We also report here new experimental results: the possible sensory receptor gene *srh–2* is expressed under conditions where L1 aggregation takes place. Mutant worms whose *srh–2* gene has been knocked out aggregate, but their aggregates are irregularly shaped, unlike the uniformly circular aggregates of wild-type worms. Our modeling suggests this phenotype could be explained by a faster-than-normal decrease in worm movement in the mutant. This observation is potentially testable by tracking the movement of individual fluorescently labeled worms.

### Validity of the continuum approximation

Two useful approaches to modeling the movements of populations are individual-based models and continuum approximations. In the individual-based model (also known as a Lagrangian model by analogy to classical mechanics) the population is represented as a collection of agents, each of which moves and changes state according to its own biological imperatives. In the continuum approximation (also known as a Eulerian model) the population is instead represented as a continuous density function of state variables, such as position. As the population is in fact composed of individuals, the individual-based model has greater *prima facie* validity. Continuum models, however, are often more tractable numerically and analytically.

We used both approaches but relied most heavily on continuum models. How valid is the continuum approximation in this case? To what extent does it distort our results? The clearest way in which continuum results (Fig 2, 3) differ from experimental results (Fig 7A) and an individual-based model (Fig S3) are the direct effect of the continuum approximation: the density function *ρ*(**x**), which is constrained to be continuous and differentiable, is smooth, while the actual distribution of worms is inevitably lumpy. This shows up in two obvious ways. First, aggregates in continuum simulations have smooth edges. The edges of the aggregates in experiments have worm-shaped irregularities. Similarly, the aggregates in individual-based simulations show irregularities shaped like simulated worms.

Second, regions of low density in individual-based simulation or experimental results are mostly empty, but with entire worms dotted here and there. Continuum simulations, in contrast, show regions of uniform low *ρ*. In these regions, however, the continuum results are not so much wrong as subject to a more subtle interpretation. That is *ρ*(**x**) is best understood as a measure. That is, for a region *R* ⊂ ℝ^*d*^, let

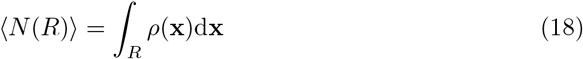

Then ⟨*N* (*R*) ⟩ is the expected number of worms in region *R*. Where ⟨*N* (*R*) ⟩ ≪ 1, it can be understood as the probability of finding one worm in region *R*.

Mogilner et al [31], in studying an individual-based Keller-Segel model state the following criterion, “A requirement for the validity of the Eulerian approximation is that many organisms are located on a spatial scale on the order of the range of interactions.” I.e., for the continuum approximation to be valid, no individual worm should matter very much. The continuum approximation is valid at most to the extent that, if we were to remove a single worm, none of the other worms would notice.

In our models the range of attractant is 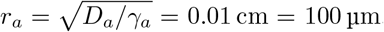. At density *ρ*, a crude estimate of the sphere of influence of a worm—the number of worms in two-dimensional space influenced by a single worm, as well as the number of worms that influence a single worm—is the number within a circle of radius *r*_*a*_ = 0.01 cm, *ρπr*^2^. Since the mean density in the full-scale simulation (Fig 6) is 2000 cm^−2^, this suggests that the sphere of influence of one worm is 0.63 worms, which is certainly not “many organisms”. This calculation, however, is obviously misleading. A glance at Fig 6 shows that the mean density is low because much of the surface is empty space. Mean density 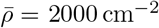 is the density weighted by surface area, i.e., it is an average in which every square millimeter of real estate counts equally. Since we are interested in the worms rather than the agar surface, we should instead compute a worm-weighted average,

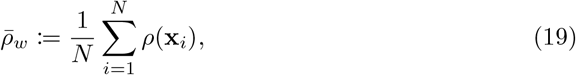

where *N* is the total number of worms and **x**_*i*_ is the location of worm *i*. The number of worms within the sphere of influence of an average worm in Fig 6 is then 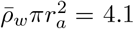.

Mogilner et al’s [31] analysis is not directly applicable to our model because they make the analytically convenient but psychophysically implausible assumption that influences are additive. This assumption doesn’t hold here because potential (7) is nonlinear. To apply their criterion we have to linearize potential around the attractant concentration the worms experience. At the worm-weighted mean attractant concentration (defined analogously to (19)) 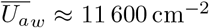, the elasticity of 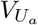 is

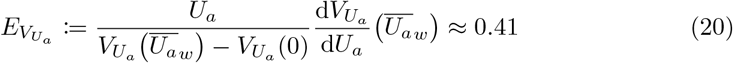

Thus, if you remove one worm, the worms in its vicinity feel a relative change in attractant potential equivalent to about 0.41/4.1 = 10% of the total attractant effect. These calculations suggest that the continuum approximation may not be drastically inaccurate.

The similarity of the results of PDE solutions (Fig 2, 3) and individual-based simulations (Fig S3) supports this conclusion. This is the more remarkable as the parameters of the PDE model and the cellular Potts model don’t correspond. As a result, cellular Potts model simulations are not expected to produce results that agree in quantitative detail to those of the PDE model, not even statistically. In addition, the cellular Potts model is defective as a model of *C elegans* chemotaxis. It would therefore be incorrect to regard the cellular Potts model as a correct model to which the PDE model is a continuum approximation. Both models are wrong, although wrong in different ways—i.e., the PDE model approximates a finite, discontinuous worm population with a continuous function *ρ*(*t*, **x**) of space and time, while the cellular Potts model models the worm and its chemotaxis in a biological unrealistic way. To the extent that the results nevertheless agree, we may be reassured that they are not the effects or idiosyncratic characteristics such as being continuum or individual-based.

## Materials and methods

### Numerical solution of partial differential equations (PDEs)

We simulated *C elegans* L1 aggregation by solving PDEs (1, 2) or (13, 14, 2) numerically in one or two spatial dimensions. In models with repellent and attractant, there were three PDEs, one for *ρ* (2) *and one each for U*_*a*_ (13), *and U*_*r*_ (14). The domain for one-dimensional simulations was a simple interval Ω = [0, *w*]. Domains for two-dimensional simulations were rectangular Ω = [0, *w*] × [0, *h*]. Width *w* and height *h* varied according to the problem. To avoid distortion of the behavior by boundary effects, all simulations were carried out with periodic boundary conditions. (For an explanation and examples of these boundary effects, see Avery [26]).

Continuous fields *ρ, U*_*a*_ and *U*_*r*_ were approximated by a grid of points equally spaced in each dimension. The spatial derivatives in the PDEs were replaced with linear combinations of the function values *ρ, U*_*a*_, *U*_*r*_, and *V* (*ρ, U*_*a*_, *U*_*r*_) (7, 9) to approximate the time derivative of each field at each point to fourth order. Simulation of the attractant+repellent model at a resolution of 384 cm^−1^ on a 6 cm × 6 cm domain requires 3 × (6 × 384)^2^ = 15 925 248 degrees of freedom.

We implemented the solution of this system of ODEs (ordinary differential equations) in PETSc (the “Portable Extensible Toolkit for Scientific computation”) [32, 33, 34]. Among the tools included in PETSc is the `TS` (time-stepper) package, a library of ODE/DAE (differential algebraic equation) solvers [35]. All solutions shown were produced with the PETSc Rosenbrock-W time stepper `ra34pw2` [36], an implicit third-order method. We used PETSc’s basic adaptive step size mechanism. This method uses error estimates from the embedded stepper to adjust step size so as to maintain error below predetermined absolute and relative tolerances. In addition, we imposed a step size limit inspired by the Courant-Friedrichs-Lewy (CFL) condition. At each step we calculated the mean worm velocity **v** = −**∇***V* at each point. We limited step size to min(|Δ*x/v*_*x*_|, |Δ*y/v*_*y*_|). (Here Δ*x* and Δ*y* are the point spacing in the *x* and *y* directions, and the minimum is taken over both dimensions and all spatial points.)

Linear equations were solved with the MUMPS parallel direct solver [37, 38] for one-dimensional problems and with PETSc’s built-in `gmres` (generalized minimal residual) Krylov solver for two-dimensional problems.

### Cellular Potts model simulations

Individual-based model simulations were run in the Morpheus [29] modeling environment, using a cellular Potts model [39, 40] to model worm movement. We sought to develop models that corresponded as closely as possible to the PDE model. PDEs describing chemical fields (1, 13, 14) can be reproduced exactly in Morpheus.

Parameters of the cellular Potts model, unfortunately, do not correspond in any simple way to those that determine worm velocity and dispersal in *ρ* PDE (2). Although valiant efforts have been made to relate the cellular Potts model to continuum models [41, 42], these require such drastic simplifying assumptions (e.g. that worms are coordinate axis-aligned rectangles) as to be practically useless. Instead, we calibrated the cellular Potts model by running it with a wide range of parameter values. Perfect calibration is not possible because, for instance, mean velocity varies nonlinearly with the strength of the chemotactic potential gradient. Although this problem can in principle be fixed by taking very small time steps, that would exacerbate an already severe computational feasibility problem. (The simulation shown in Fig S3C,D,E required 9 days. While 9 days is not an unreasonable time to wait for a result if one does everything right the first time, for such sublunary beings as ourselves a 9 day run time is a serious inconvenience.)

### Full-scale simulations

Full-scale simulations were carried out on a 6 cm × 6 cm square. The initial condition placed 72 000 worms on the square, for a final mean density 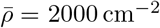. (This mean density is much lower than the mean density 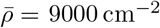 used for small-scale simulations with uniform density, because in the full-scale simulations worms are concentrated near the center of the domain, where the density is higher than the mean.) These 72 000 worms were made up of 3600 distributed uniformly on the plate to avoid zero or negative densities, which would result in (2) becoming undefined, plus 68 400 worms placed in a 2 cm diameter circle at the center of the square, with the density in the square as if a 2 cm diameter sphere had been placed in the center and the worms in it fell vertically onto the surface.

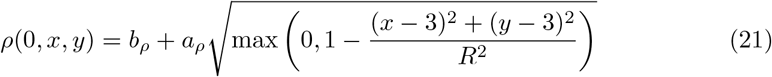

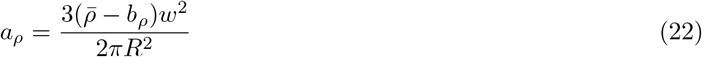

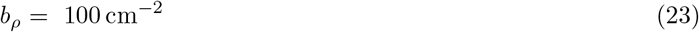

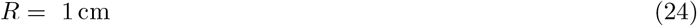

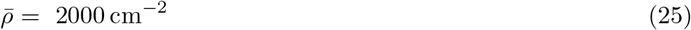

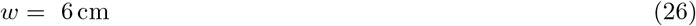

To this initial condition was added normally distributed pseudorandom noise with standard deviation 0.01*ρ*(0, *x*_*i*_, *y*_*j*_) at each grid point (*x*_*i*_, *y*_*j*_). In addition, to simulate the continuous production of noise that occurs in real experiments, pseudorandom noise was injected in the course of the simulation. Noise generation was modeled as an independent geometric Brownian motion attached to each grid point. Noise was injected at times *t*_*n*_ = 10^*n/*2^ s for *n* from 0 to 10, i.e., at 1 s, 3.16 s, 10 s, 31.6 s, 100 s, 316 s, 1000 s, 3162 s, 10 000 s, 31 622 s and 100 000 s. To inject noise, *ρ*(*t*_*n*_, *x*_*i*_, *y*_*j*_) was multiplied by a pseudorandom number exp{P_*n*_(*x*_*i*_, *y*_*j*_)*}* where P_*n*_(*x*_*i*_, *y*_*j*_) are independent normal pseudorandom variates with variance 10^−6^Δ*t*, Δ*t* being the amount of time passed since the last noise injection. After noise injection, *ρ*(*t*_*n*_, *x*_*i*_, *y*_*j*_) were normalized so that the total number of worms remained unchanged. The timing of noise injection was a pragmatic compromise. Computational efficiency precludes injecting noise continuously, since time steps had to be small immediately after noise injection—high spatial frequency noise resulted in rapid worm movement. Worms moved rapidly near the beginning of the simulation and more slowly near the end, as they approach a stable equilibrium. We thus chose a schedule in which the frequency of noise injection decreased steadily with time.

Unfortunately, numerical solution of the cellular Potts model version of the full-scale simulation was computationally infeasible.

### Spectral analysis

For spectral analysis, the final frame of the video `N2_5e5_washed.mp4` was cropped to a 960 × 960 pixel square—this corresponds to a 1.93 cm × 1.93 cm area of the agar surface. The image was standardized so that brightness *b* varied from 0 to 1, and the discrete Fourier transform calculated with Mathematica [43] function `Fourier`. This produces a two-dimensional array of Fourier coefficients 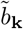, with wavenumber vector **k** = (*k*_*x*_, *k*_*y*_) *∈* (2*π/w*) ℤ^2^. Power at wavenumber **k** was computed as 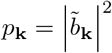. *k*_*x*_and *k*_*y*_ range from (−479) × 2*π/w* to 480 × 2*π/w*, where *w* = 1.93 cm is the width of the square. Fig 7C plots power for coefficients with *k*_*x*_/(2*π*) ranging from 0 to 20 cm^−1^ and *k*_*y*_/(2*π*) from −20 to 20 cm^−1^. We don’t show the power for *k*_*x*_ *<* 0 because the power spectrum is even in **k**, i.e. *p*_−**k**_ = *p*_**k**_. *k/*(2*π*) is the inverse of the wavelength of the corresponding sinusoid, so, for instance, *k*_*x*_/(2*π*) = 10 cm^−1^ corresponds to a sinusoid whose wavelength in the *x* direction is 1 mm.

For comparison to experiment a 200 000 s image of the full-scale simulation was cropped and scaled and standardized to a [0, 1] range. The cropping square was chosen so that the center point of the simulation was located at the same place as the center of the central mass in the cropped experimental image. The scale was chosen to make the major peaks in the radial power spectrum (Fig 7F) near 2*π×*9.5 cm^−1^ correspond precisely. Finally a black circle of the same dimensions as the central mass in the video image was added.

The radially summed power spectrum was approximated by an array *s* of 1024 numbers representing power at wavenumbers from 0 to 2*π ×* 20 cm^−1^. Each wavenumber in the two-dimensional power spectrum was mapped linearly to a point in *s*. Since most wavenumbers mapped to noninteger locations, they were represented by a weighted sum of the two closest elements of *s*. For instance, consider **k** = 2*π ×* (1, 2)/1.93. This corresponds in the radial spectrum to wavenumber 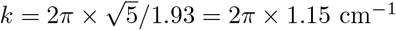. This *k* maps to radial array *s* location *j* = 1 + 1023(1.15/20) = 60.26. If *p*_**k**_ is the power at **k** in the two-dimensional spectrum, we add (61 − *j*)*p*_**k**_ to *s*[60] and (*j* − 60)*p*_**k**_ to *s*[61].

The radially summed spectrum thus computed is noisy and quasi-periodic with period 1023/(20 × 1.93) ≈ 26.5, reflecting the discrete two-dimensional power spectrum. To produce Fig 7F, we smoothed *s* using Mathematica function `GaussianFilter` with smoothing radius 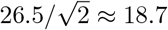 which we determined by trial and error to be the smallest smoothing radius that effectively eliminated the periodic structure.

### Effect of ethanol and acetate on the transcriptome of starved L1s

To obtain L1s for transcriptomic profiling, we grew N2 worms in liquid culture. We inoculated 250 mL S-complete in a 2 L flask with 7 × 10^5^ synchronized L1 larvae obtained from a small-scale liquid culture and added 10 mL 50% *E. coli* K-12 stock suspension. Worms were grown at 22°C, 220 rpm (for details see Artyukhin et al [2]). We monitored the worm culture during the next 2.5 days and added *E. coli* food as it became depleted. Bleaching of gravid adult worms (8 mL water + 2 mL bleach + 0.3 mL 10 M NaOH for 6 min) after 68 h of growth yielded ca. 1 × 10^7^ eggs. After 3 washes with M9 buffer, eggs were resuspended in 20 mL M9 and allowed to hatch. After 2 days of starvation (51 h to 53 h after bleaching, 20°C), we collected L1 larvae, washed them 6 times with M9, 10 mL each time, and resuspended in 10 mL M9 after the final wash. 300 µL of the resulting L1 suspension (corresponding to ca. 20 µL L1 pellet) were added to each of the following solutions in 15-ml plastic tubes: (1) 3 mL M9; (2) 3 mL M9 + 3 µL ethanol; (3) 3 mL M9 + 51 µL 1 M potassium acetate. Tubes were put on a rocker at room temperature (ca. 21°C). After 1.5 h, the tubes were cooled down on ice for 1 min, centrifuged, and ca. 2 mL of supernatant was removed by aspiration. We resuspended L1s in the remaining ca. 1 mL of liquid, transferred the suspension to mL tubes, centrifuged, removed supernatant, added 300 µL Trizol to each sample, and froze in liquid nitrogen. The above procedure was repeated two more times with worms grown on different days to obtain biological triplicates for each condition (control, ethanol, potassium acetate). All samples were stored at -80°C before analysis.

### Microarray methods

Microarray methods were as described by Hyun et al [44].

### *srh–2* knockout

Deletion mutants of *srh–2* were generated by CRISPR and obtained from Knudra. Three deletion strains were generated (COP-1274, 1275, 1276), all of the same genotype: *unc–119(ed3) III; srh–2(knu317::unc–119(+)) V*.

## Acknowledgements

We thank Leah Edelstein-Keshet for introducing us to the Morpheus modeling environment.

## Supporting information

### Linear stability analysis of the attractant-only model

The attractant–only model (1, 2) shares with the original Keller-Segel system [21] the property of density-dependent instability. There is a uniform equilibrium,

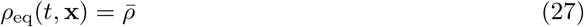

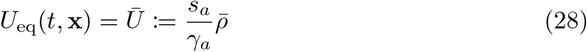

Substituting these functions into (1, 2) shows that d*ρ*_eq_/d*t* = d*U*_eq_/d*t* = 0. Consider a population near this equilibrium, and 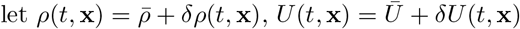, *U* (*t*, **x**) = *Ū* + *δU* (*t*, **x**). Because 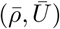 is an equilibrium, *ρ*_*t*_(*t*, **x**) = *δρ*_*t*_(*t*, **x**) and *U*_*t*_(*t*, **x**) = *δU*_*t*_(*t*, **x**).

Substituting into (1, 2),

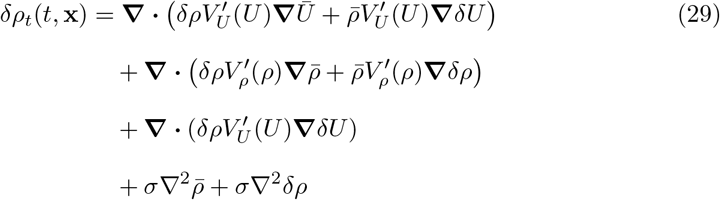

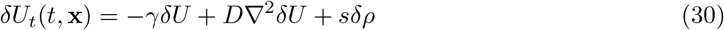

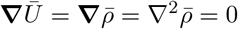.Writing 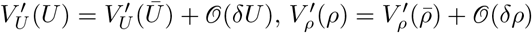 and ignoring second order terms we have, to first order, the linear vector PDE,

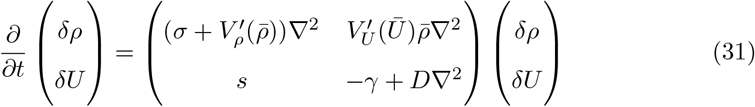

This is easily solved by separation of variables. It will be convenient to define 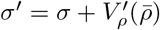. Since *V*_*ρ*_ is, by design, an increasing function, 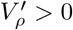 and *σ ′ > σ >* 0. For the parameters in Table 2, 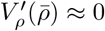 and *σ ′* ≈ *σ*. Now, eigenfunctions of (31) are of the form

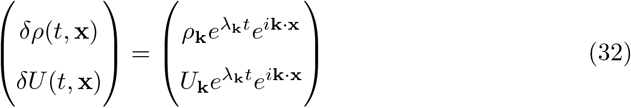

**k** is a wavenumber vector, i.e., a vector of frequency in each spatial direction. Substituting into (31) produces the 2 × 2 matrix eigenvalue problem,

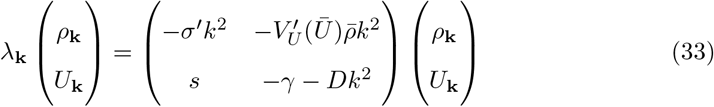

where *k*^2^ := ‖ **k** ‖ ^2^. Solutions of the linearized system (33) are linear combinations of functions (32) where (*ρ*_**k**_, *U*_**k**_)^T^ is an eigenvector of the matrix in (33). If, for every **k**, Re(*λ*_**k**_) *<* 0 then any small fluctuation away from the uniform equilibrium will die away, and the uniform equilibrium will be stable. If, however, there exists a **k** such that the matrix has an eigenvalue with positive real part, then the uniform equilibrium is unstable. The sum of the two eigenvalues, the trace of the matrix, is negative,−*σ ′ k*^2^ − *γ* − *Dk*^2^ *<* 0, so the only possible way to have an eigenvalue with positive real part is if both eigenvalues are real, one positive and one negative. Thus, first-order instability is expected if and only if the determinant is negative,

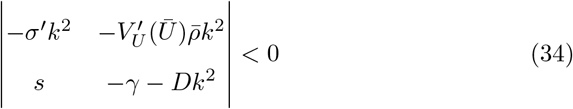

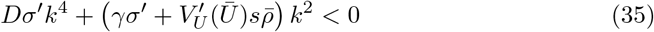

Inequality (35) can hold only if 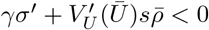. Since, as mentioned above, *V*_*U*_ is a decreasing function of attractant concentration, 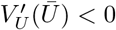. If the condition 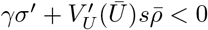 holds, then for some small enough *k*^2^, the negative *k*^2^ term will dominate the positive *k*^4^ term and the determinant will be negative. Thus, first-order instability occurs if and only if 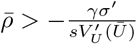. Since 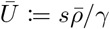 depends on 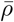, it is not a foregone conclusion that instability is possible. Neglecting 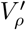 and with *V*_*U*_ as in (7), the instability condition reduces to

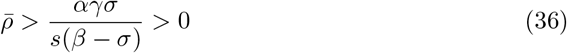

Instability is possible if *β > σ*. By design, the instability condition is 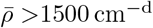 with the parameter values in Table 2.

### Linear stability analysis of the attractant+repellent model

For simplicity, we assume *V*_*ρ*_ = 0 in the following analysis. With the parameter values in Table 2, this is very close to true in the vicinity of the instability threshold. Linearization of the PDE system (2, 13, 14) around a uniform equilibrium at 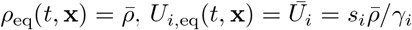 produces the following linear PDE system

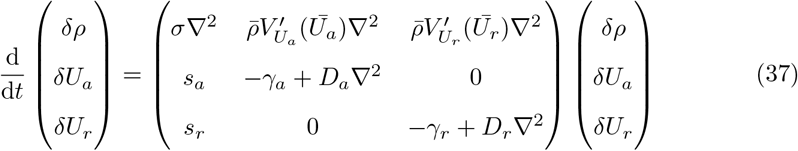

The ansatz

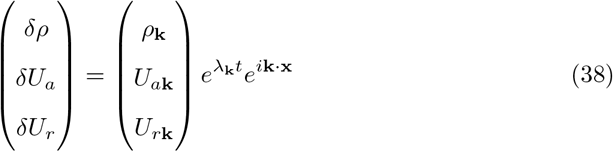

yields the eigenvalue problem

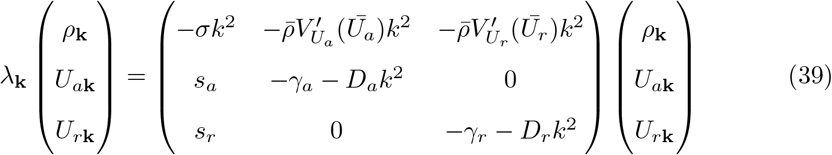

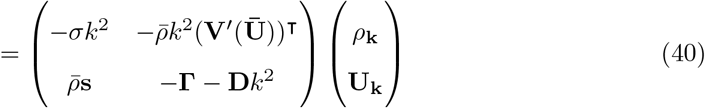

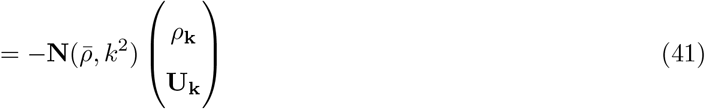

In (40), the matrix is in a block form that can be extended easily to any number of signals, with

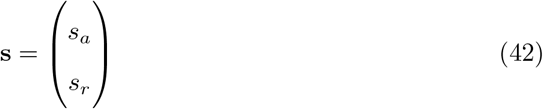

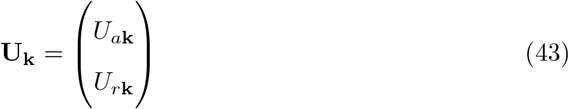

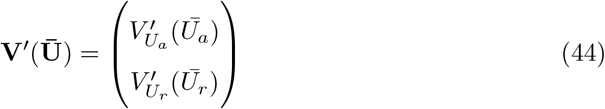

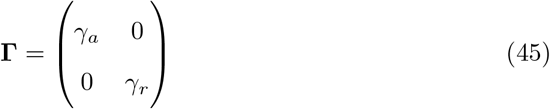

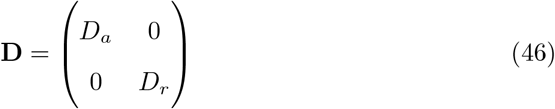

In (41), 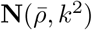 is defined as the negative of the matrix in (40). (We define **N** as the negative to avoid an inconvenient factor of 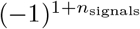 in the determinant we are about to calculate.) The uniform equilibrium is unstable at mean density 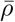 if for some 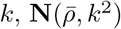 has an eigenvalue with negative real part. If 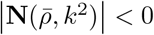, the equilibrium is certainly unstable. This leads to the following criterion for instability

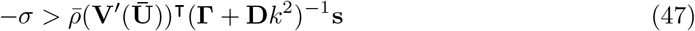

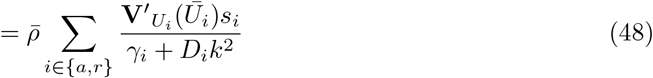

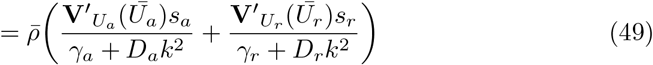

It is possible to choose parameter values so that this criterion predicts instability with a nontrivial minimum wavenumber (and therefore finite maximum scale). How does this work? Remember that 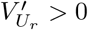 because it is a repellent and 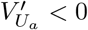 because it is an attractant. Thus the two terms in (49) are opposite in sign. Also, *γ*_*r*_ *< γ*_*a*_ and *D*_*r*_ *> D*_*a*_, because the repellent is a longer-range signal than the attractant. Thus, at low *k*, if the relative magnitudes of 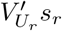 and 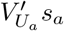 are appropriately adjusted (by evolution or the modeler), the sum in (49) is positive and the uniform equilibrium is stable to perturbations of small wavenumber = large scale. As *k* rises the *D*_*r*_*k*^2^ factor in the denominator of the repellent term makes the positive term small compared to the negative attractant term. The sum in parentheses can become negative, and if 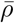 is large enough, the right-hand-side drops below −*σ*, and instability to perturbations of intermediate wavenumber = medium scale results. For large *k* the right-hand-side approaches zero because of the *Dk*^2^ factors in both denominators. The uniform equilibrium is thus stable to perturbations of large wavenumber = small scale. It is therefore possible for an attractant+repellent Keller-Segel model to have a finite natural scale.

Based on calculations of this sort we chose *β*_*r*_ = −*β*_*a*_ = −2*σ* = −5.56 × 10^−4^ cm^2^s^−1^. Attractant parameters remained as in Table 2. The addition of a repellent increases the threshold for instability, but the uniform equilibrium is still predicted to be unstable at 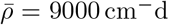.

### Convergence tests

To test whether the numerical solution of the worm system PDEs (2, 13, 14) approximates the correct solution, we compared numerical solutions of the attractant+repellent system to an analytical solution of the linearized PDE system (37).

In one dimension, for *x ∈* Ω = [0, 1],

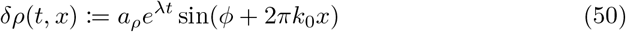

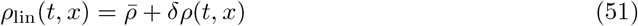

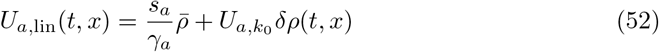

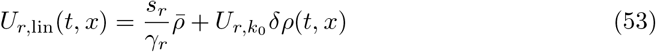

In two dimensions, for 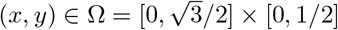,

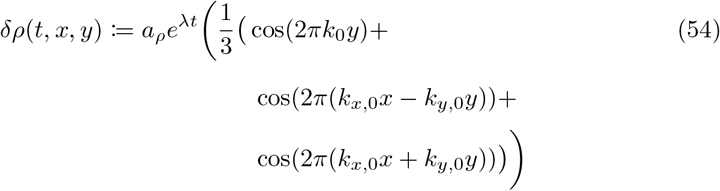

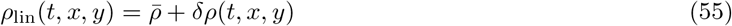

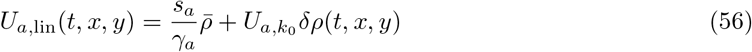

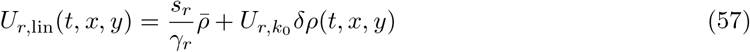

Here *ϕ, a*_*ρ*_ ∈ ℝ and *k*_0_ ∈ 2 ℤ are parameters that can be freely chosen. We chose *a*_*ρ*_ = 1 and *ϕ* = *π/*2. We chose *k*_0_ = 4 to produce a substantial positive growth rate. For the two-dimensional case, we chose 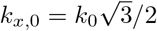 and *k*_*y*,0_ = *k*_0_/2 to produce hexagonal symmetry. *λ* is the positive eigenvalue of 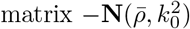 (41), with corresponding eigenvector 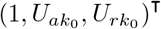 (as in (38), but normalized so that 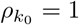). Numerical values *λ* ≈ 0.000 955 s^−1^, 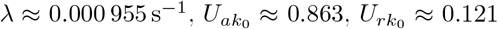 were estimated to 15-digit precision by numerical diagonalization of the computed 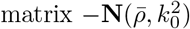.

Functions (*ρ*_lin_(*t, x*), *U*_*a*,lin_(*t, x*), *U*_*r*,lin_(*t, x*))^T^ are of course not an exact solution of the full nonlinear PDEs (2, 13, 14). To produce a closely related system with this exact solution for convergence testing, we modified the *ρ* PDE (2) by addition of a source term *S*(*t*, **x**).

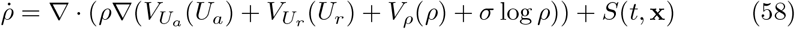

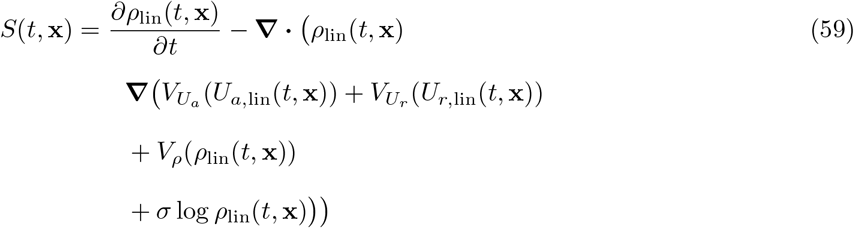

It was unnecessary to modify the *U*_*a*_ and *U*_*r*_ PDEs since they are linear. Source function (59) was computed symbolically from linear solutions (51-53) or (55-57) and converted to `sympy` expressions with Mathematica [43]. `sympy` is a computer algebra package for the programming language `python`.

### Software

Software developed for this work is available from https://github.com/leonavery/KSFD. Morpheus [29] implementations of the cellular Potts models are available from https://github.com/leonavery/worm-CPM. The README file for worm-CPM discusses the suitability of the cellular Potts model for worms in some detail.

## Supplemental figures

### Supplemental data

The following data file is provided:

**C**.**elegans microarray results_020615_CID**.**xlsx** Microarray expression profiling results.

### Videos

The following video files are provided.

Avery, Leon (2021), “Avery_L1agg2”, Mendeley Data, V3, doi: 10.17632/r5v772ftcs.3

**Table S1.**
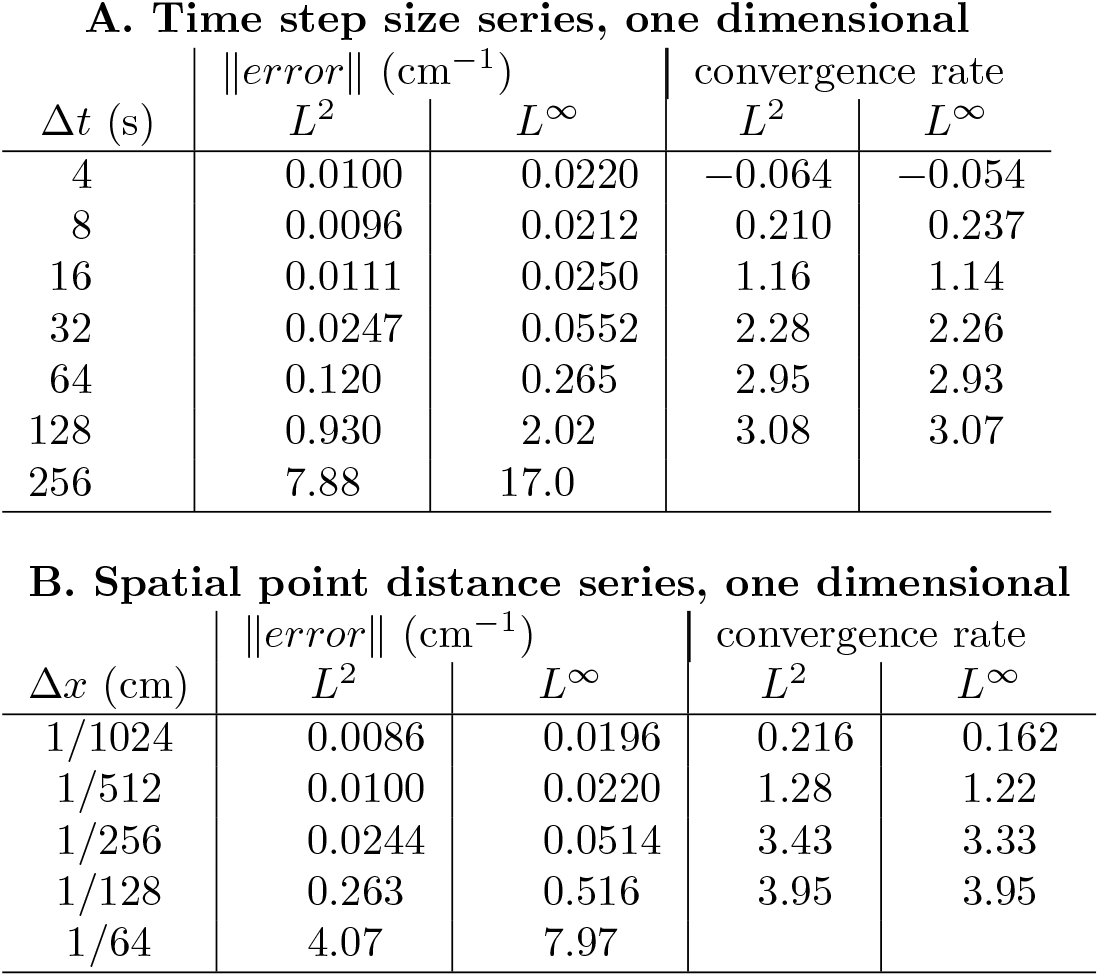
Convergence test results, one spatial dimensional Eqs. (58, 13, 14) were solved numerically from *t* = 0 s to 8192 s on *x∈* Ω = [0, 1]. In this time the amplitude of the sinusoid *δρ* (50) *grew from a*_*ρ*_ = 1 to *a*_*ρ*_*e*^8192*λ*^ ≈2505. **A** shows the results of varying the time step size from 4 to 256 s (with a fixed spatial point distance of Δ*x* =1/512 cm). **B** shows the results of varying the spatial point distance from 1/1024 to 1/64 cm (with a fixed time step of 4 s). The error in *ρ* at the final time point was calculated as the difference between the numerical result and exact result (51). *L*^2^ and *L*^*∞*^ norms of the error are tabulated. Convergence rate is calculated between consecutive rows as log(‖error_1_ ‖*/ ‖* error_2_ ‖)/ log(*h*_1_*/h*_2_), with *h* being either Δ*t* or Δ*x*, as appropriate. The mean of *ρ* was 9000 cm^−1^ in all cases. Thus the relative error is about 1/9000 times the error shown—e.g. 0.0100/9000 ≈ 1.1 × 10^−6^ for Δ*t* = 4 s, Δ*x* = 1/512 cm in one dimension. Errors in *U*_*a*_ and *U*_*r*_ (not shown) were smaller but otherwise behaved similarly. All numerical solutions used the PETSc Rosenbrock-W solver `ra34pw2` (nominally an order 3 method), and fourth-order approximations for the spatial derivatives.

A. Time step size series, one dimensional
| $\Delta t$ (s) | $\ error\ $ (cm<sup>-1</sup>) | | convergence rate | |
| --- | --- | --- | --- | --- |
| | $L^2$ | $L^\infty$ | $L^2$ | $L^\infty$ |
| 4 | 0.0100 | 0.0220 | -0.064 | -0.054 |
| 8 | 0.0096 | 0.0212 | 0.210 | 0.237 |
| 16 | 0.0111 | 0.0250 | 1.16 | 1.14 |
| 32 | 0.0247 | 0.0552 | 2.28 | 2.26 |
| 64 | 0.120 | 0.265 | 2.95 | 2.93 |
| 128 | 0.930 | 2.02 | 3.08 | 3.07 |
| 256 | 7.88 | 17.0 |  |  |

B. Spatial point distance series, one dimensional
| $\Delta x$ (cm) | $\ error\ $ (cm<sup>-1</sup>) | | convergence rate | |
| --- | --- | --- | --- | --- |
| | $L^2$ | $L^\infty$ | $L^2$ | $L^\infty$ |
| 1/1024 | 0.0086 | 0.0196 | 0.216 | 0.162 |
| 1/512 | 0.0100 | 0.0220 | 1.28 | 1.22 |
| 1/256 | 0.0244 | 0.0514 | 3.43 | 3.33 |
| 1/128 | 0.263 | 0.516 | 3.95 | 3.95 |
| 1/64 | 4.07 | 7.97 |  |  |

**Table S2.** Convergence test results, two spatial dimensions Eqs. (58, 13, 14) were solved numerically from *t* = 0 s to 8192 s on 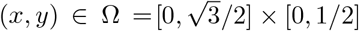. In this time the amplitude of the sinusoid *δρ* (54) *grew from a*_*ρ*_ = 1 to *a*_*ρ*_*e*^8192*λ*^ ≈ 2505. **A** shows the results of varying the time step size from 4 to 256 s (with a fixed spatial point distance of 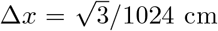). **B** shows the results of varying the spatial point distance from 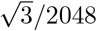 to 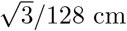 (with a fixed time step of 4 s). In all cases 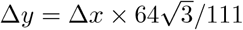. Errors and convergence rates calculated as in Table S1.

**A. Time step size series, two dimensional**
| $\Delta t$ (s) | $\ error\ $ (cm<sup>-1</sup>) | | convergence rate | |
| --- | --- | --- | --- | --- |
| | $L^2$ | $L^\infty$ | $L^2$ | $L^\infty$ |
| 4 | 0.0072 | 0.0469 | 2.01 | 2.20 |
| 8 | 0.0290 | 0.215 | 0.692 | 0.627 |
| 16 | 0.0468 | 0.332 | -1.57 | -2.21 |
| 32 | 0.0158 | 0.0716 | 3.40 | 3.48 |
| 64 | 0.167 | 0.800 | 1.53 | 1.28 |
| 128 | 0.483 | 1.94 | 3.14 | 3.21 |
| 256 | 4.27 | 18.0 |  |  |

| $\Delta x$ (cm) | $\ error\ $ (cm<sup>-1</sup>) | | convergence rate | |
| --- | --- | --- | --- | --- |
| | $L^2$ | $L^\infty$ | $L^2$ | $L^\infty$ |
| $\sqrt{3}/2048$ | 0.0246 | 0.178 | -1.77 | -1.92 |
| $\sqrt{3}/1024$ | 0.0072 | 0.0469 | 0.103 | -0.172 |
| $\sqrt{3}/512$ | 0.0077 | 0.0416 | 2.84 | 2.26 |
| $\sqrt{3}/256$ | 0.0555 | 0.199 | 3.92 | 3.86 |
| $\sqrt{3}/128$ | 0.838 | 2.88 | | |

**Figure S1.**
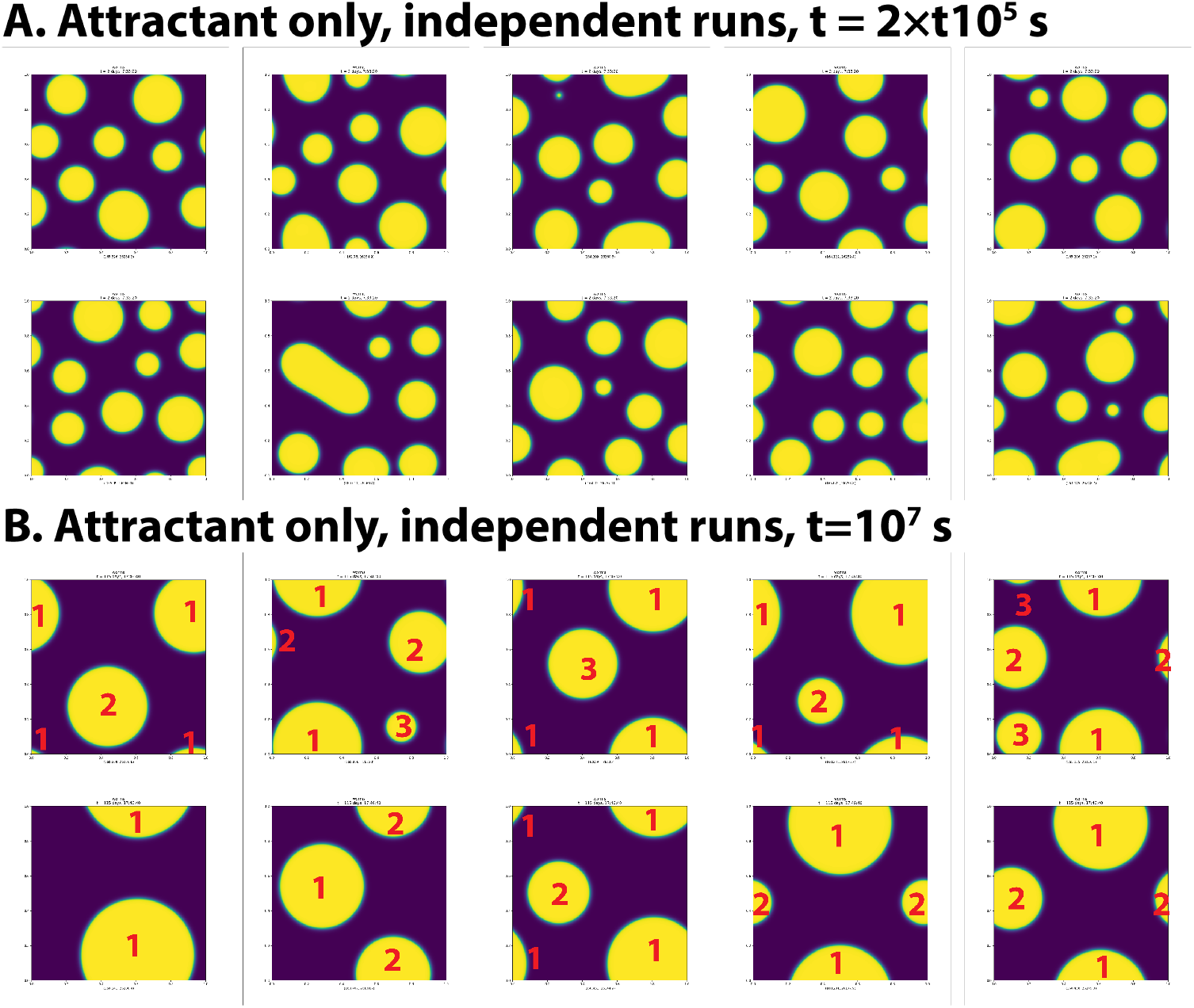
Attractant-only simulation reruns **A**. These ten images reproduce the numerical experiment of Figure 2B—simulation of the attractant-only model in two dimensions—but with different pseudorandom noise in the initial condition. Only worm density *ρ*(*x, y*) at *t* = 200 000 s (2 days, 7:33:20) is shown. **B**. Like **A**, but at *t* = 1 × 10^7^ s (115 days, 17:46:40). These images correspond one-to-one to the images in **A**. The number of aggregates in these panels ranges from one to three, although a single aggregate may appear in as many as four pieces because of the periodic boundary conditions. To ease the identification of aggregates, the aggregate to which each piece belongs is identified by a red number.

#### N2 5e5 washed.avi

This video shows the time course of aggregation after 500 000 starved L1s were pipetted onto the center of a petri plate. The recording covers 720 min. There is one frame per minute of real time, and the playback rate is 7 s^−1^.

#### options138a.mp4

Numerical solution of attractant-only model in one dimension.This video corresponds to Fig 2A. This and all following videos are 200s long at 15 s^−1^. Time is displayed as “days, H:MM::SS”. Time ranges from 0 s to 200 000 s (2 days, 7:33:20). The two numbers below each panel are the minimum and maximum of the plotted field.

**Figure S2.**
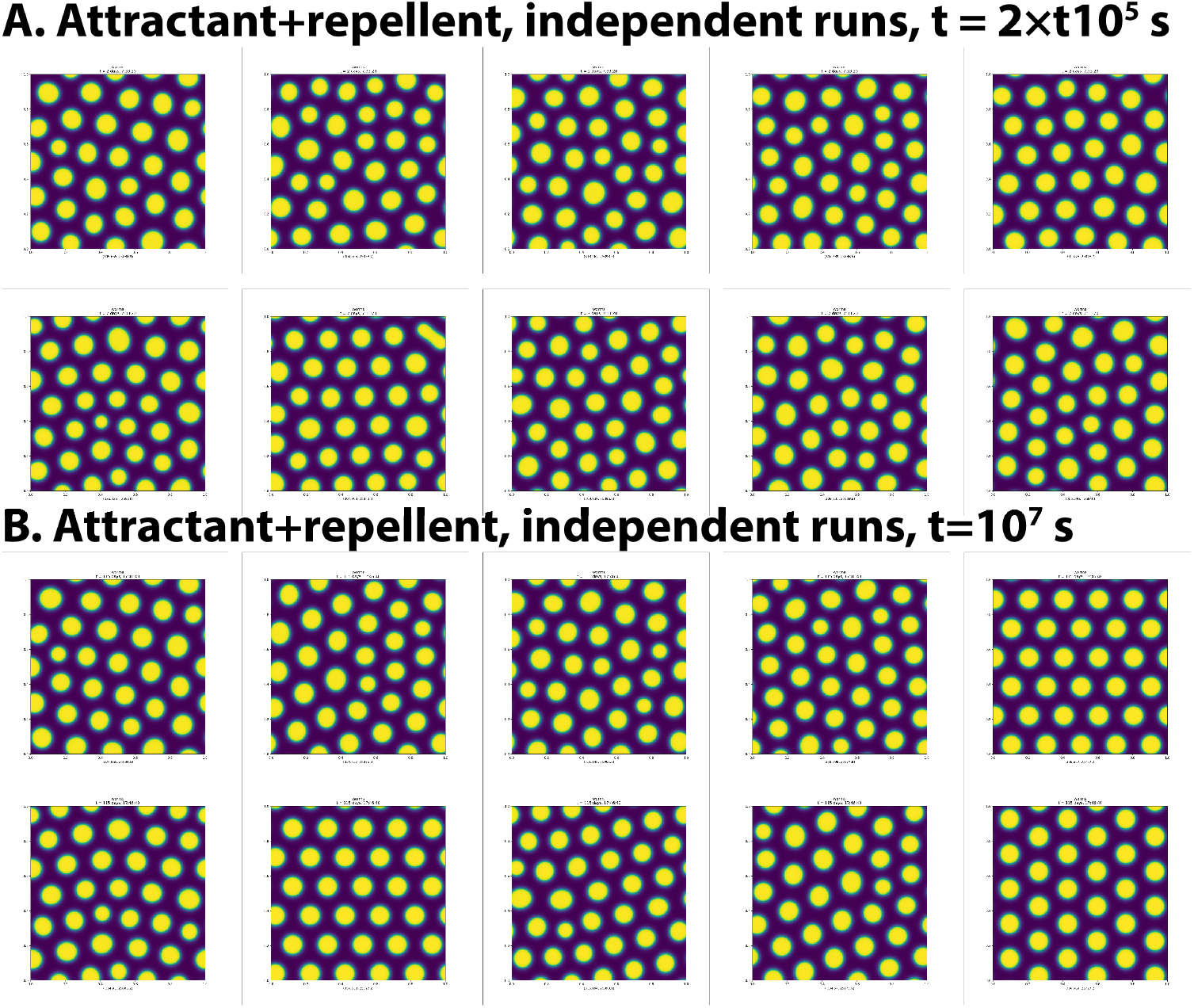
Attractant+repellent simulation reruns **A**. These ten images reproduce the numerical experiment of Fig 3B—simulation of the attractant+repellent model in two dimensions—but with different pseudorandom noise in the initial condition. Only worm density *ρ*(*x, y*) at *t* = 200 000 s (2 days, 7:33:20) is shown. **B**. Like **A**, but at *t* = 1 × 10^7^ s (115 days, 17:46:40). These images correspond one-to-one to the images in **A**.

#### options139.mp4

Numerical solution of attractant-only model in two dimensions.Corresponds to Fig 2B.

#### options140a.mp4

Numerical solution of attractant+repellent model in one dimension.Corresponds to Fig 3A.

#### options141.mp4

Numerical solution of attractant+repellent model in two dimensions.Corresponds to Fig 3B.

#### options157.mp4

Numerical solution of attractant+repellent model in two dimensions on a 6 cm × 6 cm domain, with most worms initially placed in the center.Corresponds to Fig 6.

**Figure S3.**
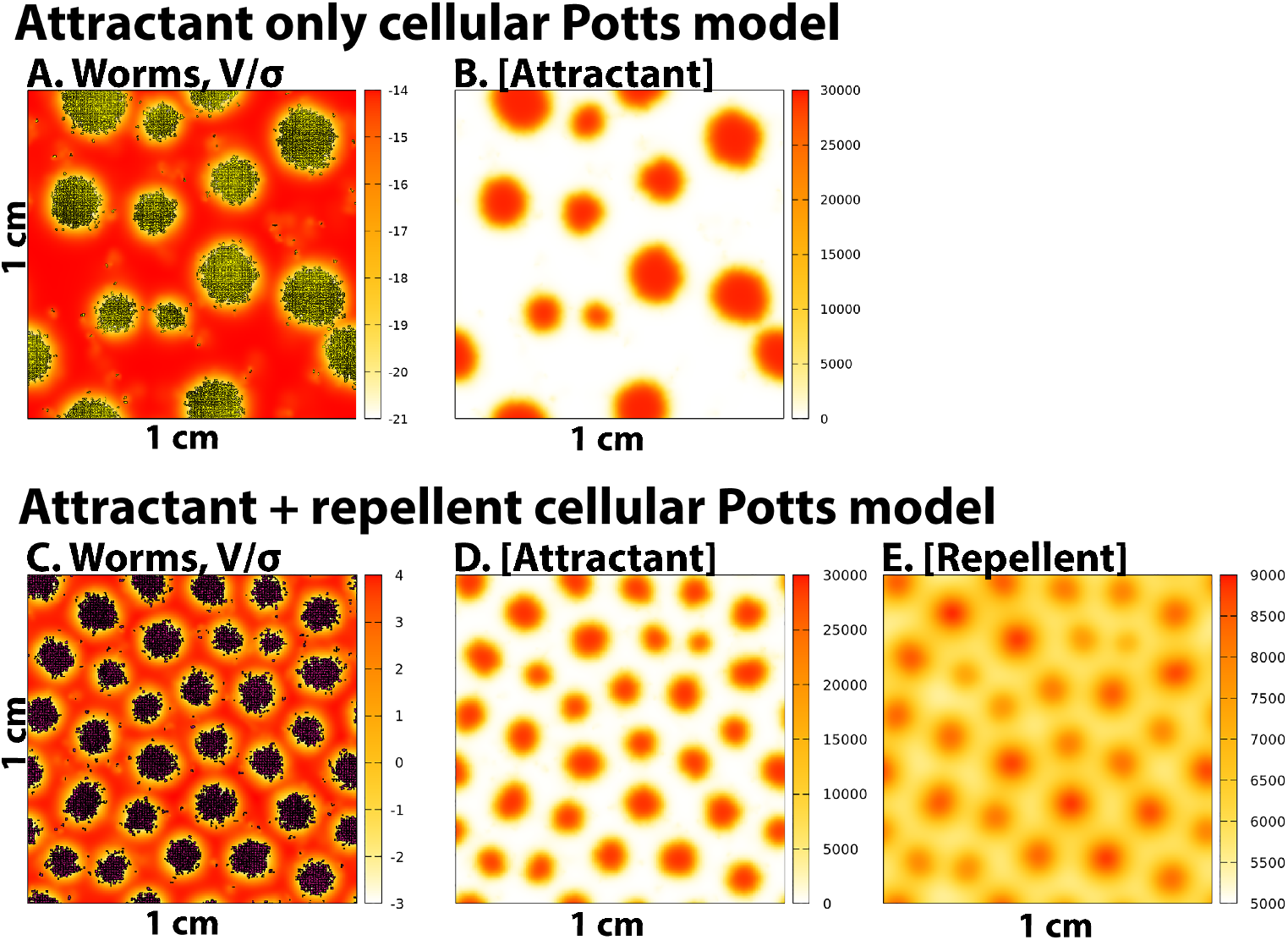
Cellular Potts model simulations **A, B** These two images show the results of an individual-based cellular Potts model simulation of the Attractant-only model in two dimensions, and are meant to be compared to Fig 2C,D. **D-F** show the results of a cellular Potts model simulation of the attractant+repellent model in two dimensions and can be compared to Fig 3C,D,E. Because there is no simple relationship between the parameters of the PDE model and the cellular Potts model, we do not expect precise quantitative agreement, even on a statistical basis.

#### worm5g.mp4

Numerical solution of attractant-only cellular Potts model in two dimensions.Corresponds to Fig S3A,B.

#### worm6c.mp4

Numerical solution of attractant+repellent cellular Potts model in two dimensions.Corresponds to Fig S3C,D,E.

